# Transcription factors interact with RNA to regulate genes

**DOI:** 10.1101/2022.09.27.509776

**Authors:** Ozgur Oksuz, Jonathan E Henninger, Robert Warneford-Thomson, Ming M Zheng, Hailey Erb, Kalon J Overholt, Susana Wilson Hawken, Salman F Banani, Richard Lauman, Adrienne Vancura, Anne L Robertson, Nancy M Hannett, Tong I Lee, Leonard I. Zon, Roberto Bonasio, Richard A. Young

## Abstract

Transcription factors (TFs) orchestrate the gene expression programs that define each cell’s identity. The canonical TF accomplishes this with two domains, one that binds specific DNA sequences and the other that binds protein coactivators or corepressors. We find that at least half of TFs also bind RNA, doing so through a previously unrecognized domain with sequence and functional features analogous to the arginine-rich motif of the HIV transcriptional activator Tat. RNA-binding contributes to TF function by promoting the dynamic association between DNA, RNA and TF on chromatin. TF-RNA interactions are a conserved feature essential for vertebrate development and disrupted in disease. We propose that the ability to bind DNA, RNA and protein is a general property of many TFs and is fundamental to their gene regulatory function.

## Main

Transcription factors (TFs), which are encoded by ∼1,600 genes in the human genome, comprise the single largest protein family in mammals. Each cell type expresses approximately 150-400 TFs, which together control the gene expression program of the cell (Lambert et al., 2018; Vaquerizas et al., 2009). TFs typically contain DNA-binding domains that recognize specific sequences and multiple TFs collectively bind to enhancers and promoter-proximal regions of genes (Avsec et al., 2021; Panne et al., 2007). The DNA-binding domains form stable structures whose conserved features are reliably detected by homology and are therefore used to classify TFs (e.g. C2H2 zinc finger, homeodomain, bHLH, bZIP) (Figure 1A) (Lambert et al., 2018; Vaquerizas et al., 2009). TFs also contain effector domains that exhibit less sequence conservation and sample many transient structures that enable multivalent protein interactions (Arnold et al., 2018; Boija et al., 2018; Soto et al., 2022). These effector domains recruit coactivator or corepressor proteins, which contribute to gene regulation through mechanisms that include mobilizing nucleosomes, modifying chromatin-associated proteins, influencing genome architecture, recruiting transcription apparatus and controlling aspects of transcription initiation and elongation (Richter et al., 2022; Vos, 2021). This canonical view of TFs that function with two domains, one binding DNA and the other protein, has been foundational for models of gene regulation (Lelli et al., 2012; Spitz and Furlong, 2012).

**Figure 1.**
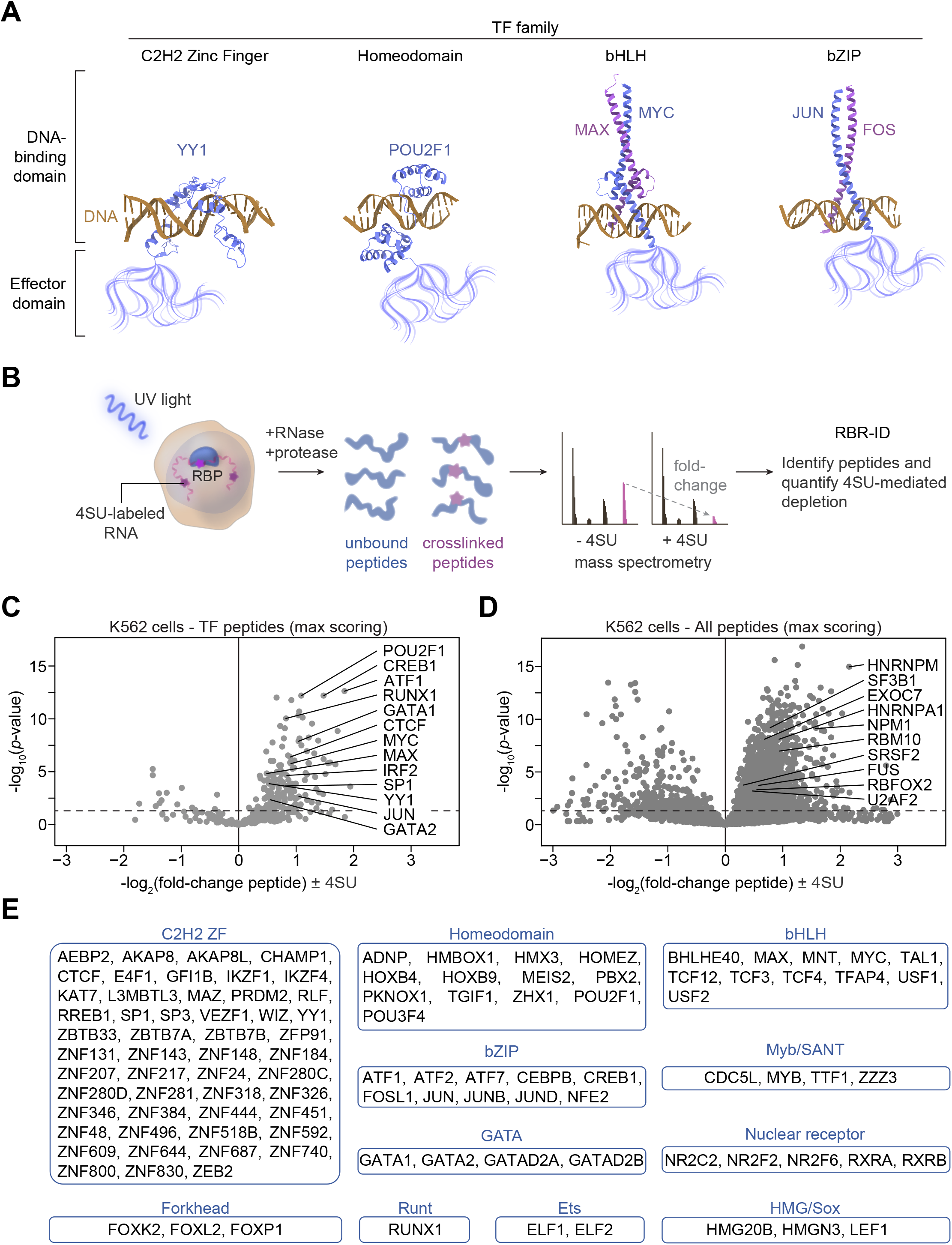
Transcription factor binding to RNA in cells. **(A)** Schematic of functional domains in transcription factors (PDB accession numbers in Methods). **(B)** Experimental scheme for RBR-ID in human K562 cells. **(C)** Volcano plot of TF peptides in RBR-ID for human K562 cells with select highlighted TFs (dotted line at p=0.05). Each marker represents the peptide with maximum RBR-ID score for each protein. **(D)** Volcano plot of all detected peptides in RBR-ID for human K562 cells with select highlighted RBPs (dotted line at p=0.05). Each marker represents the peptide with maximum RBR-ID score for each protein. **(E)** List of RBR-ID^+^ TFs (p < 0.05, log_2_FC > 0) categorized by DBD family.

RNA molecules are produced at loci where TFs are bound, but their roles in gene regulation are not well-understood (Kaikkonen and Adelman, 2018; Seila et al., 2008). A few TFs and cofactors have been reported to bind RNA (Cassiday and Maher, 2002; Holmes et al., 2020; Saldaña-Meyer et al., 2019; Sigova et al., 2015; Theunissen et al., 1992; Xu et al., 2021), but TFs do not harbor domains characteristic of well-studied RNA binding proteins. We wondered whether TFs might have evolved to interact with RNA molecules that are pervasively present at gene regulatory regions but harbor a heretofore unrecognized RNA-binding domain. Here we present evidence that a broad spectrum of TFs do bind RNA molecules, that TFs accomplish this with a domain analogous to the RNA-binding arginine-rich motif of the HIV Tat transactivator, and that this domain promotes TF occupancy at regulatory loci. These domains are a conserved feature important for vertebrate development, and they are disrupted in cancer and developmental disorders.

### Transcription factor binding to RNA in cells

Using human K562 cells, we performed a high throughput RNA-protein crosslinking assay (RNA-binding region identification - RBR-ID), which uses UV crosslinking and mass spectrometry to detect direct interactions between protein and RNA molecules in cells (Figure 1B) (He et al., 2016). The results included the expected distribution of peptides from known RNA-binding proteins (RBPs) and revealed that a broad distribution of TFs had peptides crosslinked to RNA in this assay (Figures 1C and 1D). Nearly half (48%) of TFs identified in the RBR-ID dataset showed evidence of RNA-binding in K562 cells (Figure S1A) when the analysis was conducted using thresholds that retain RBPs verified by independent methods (He et al., 2016). These results prompted a re-examination of previously published RBR-ID data for murine embryonic stem cells (ESCs) (He et al., 2016) which confirmed that a substantial fraction of TFs (41%) in those cells also bind RNA (Figures S1B-1D).

Specific TFs are notable for their roles in control of cell identity and have been subjected to more extensive study than others. Many well-studied TFs that contribute to the control of cell identity were observed among the TFs that showed evidence of RNA binding. In K562 hematopoietic cells, these included GATA1, GATA2, and RUNX1, which play major roles in regulation of hematopoietic cell genes (Orkin and Zon, 2008), as well as MYC and MAX, oncogenic regulators of these tumor cells (Delgado et al., 1995) (Figure 1C). In the ESCs, these included the master pluripotency regulators Oct4, Klf4, and Nanog, as well as the MYC family member that is key to proliferation of these cells, Mycn (Young, 2011) (Figure S1B). The RNA-binding TFs also included those involved in other important cellular processes, including regulation of chromatin structure (CTCF, YY1) and response to signaling (CREB1, IRF2, ATF1) (Figure 1C). It was notable that RNA binding was a property of TFs that span many TF families (Figure 1E and Figure S1E). These results suggest that RNA binding is a property shared by TFs that participate in diverse cellular processes and that possess diverse DNA-binding domains.

### Transcription factor binding to RNA in vitro

To corroborate evidence that TFs can bind RNA molecules in cells, we sought to confirm that purified TFs bind RNA molecules in vitro using a fluorescence polarization assay (Figure 2A, Methods). The assay was validated with multiple control proteins with an RNA of random sequence, including three well-studied RNA-binding proteins (U2AF2, HNRNPA1, and SRSF2) and proteins that were not expected to have substantial affinity for RNA (GFP and the DNA-binding restriction enzyme BamHI). The RBPs bound RNA with nanomolar affinities, consistent with previous studies (Burd and Dreyfuss, 1994; Corley et al., 2020; Maji et al., 2020; Zhang et al., 2015), whereas GFP and BamHI showed little affinity for RNA (K_d_ > 4 μM) (Figure 2B). We then selected 13 TFs that showed evidence of crosslinking to RNA in cells, are well-studied for their diverse cellular functions and are members of different TF families, purified them from human cells and measured their RNA-binding affinities. These TFs exhibited a range of binding affinities for the RNA, ranging from 41 to 505 nM, which is remarkably similar to the range of affinities measured for known RBPs (42 to 572 nM) (Figure 2C). Thus, a diverse set of transcription factors can bind RNA with affinities similar to proteins with known physiological roles in RNA processing. The thousands of enhancer and promoter-proximal regions where TFs bind have diverse sequences, and thus RNA molecules produced from these sites differ in sequence, so we investigated whether TFs bind diverse RNA sequences. Six TFs were investigated, and the results indicate that these TFs do bind various RNA sequences with similar affinities (Figure S2).

**Figure 2.**
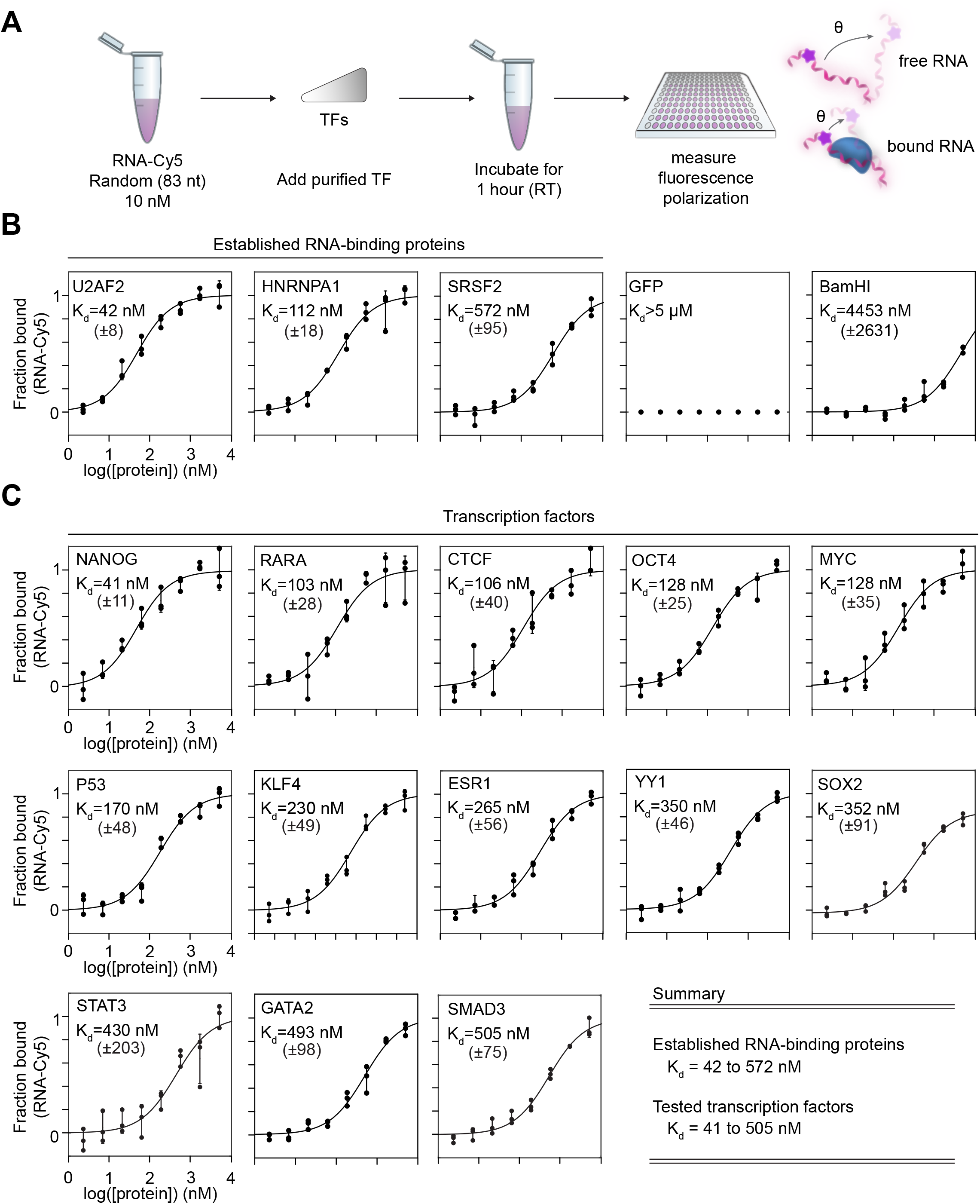
Transcription factor binding to RNA in vitro. **(A)** Experimental scheme for measuring the equilibrium dissociation constant (K_d_) for protein-RNA binding. **(B)** Fraction bound RNA with increasing protein concentration for established RNA-binding proteins, GFP, and the restriction enzyme BamHI (error bars depict s.d.). **(C)** Fraction bound RNA with increasing protein concentration for select transcription factors (error bars depict s.d.).

### An arginine-rich domain in transcription factors

We next sought to identify regions in TFs that contribute to RNA-binding. TFs do not contain sequence motifs that resemble those of structured RNA-binding domains (Corley et al., 2020; Lunde et al., 2007) (Figure S3), so we searched for local amino acid features that might be common to TFs. Nearly 80% of TFs were found to have a cluster of basic residues (R/K) adjacent to their DNA-binding domain (Figure 3A). Derivation of a position-weight matrix from these “basic patches” revealed that they contain a sequence motif similar to the RNA-binding domain of the HIV Tat transactivator, which has been termed the arginine-rich motif (ARM) (Calnan et al., 1991a, 1991b) (Figure 3B). These ARM-like domains were enriched in TFs compared to the remainder of the proteome (Figure 3C). Furthermore, the ARM-like domains have sequences that are evolutionarily conserved and appear adjacent to diverse types of DNA-binding domains, as illustrated for KLF4, SOX2, and GATA2 (Figure 3D). This analysis suggests that TFs often contain conserved ARM-like domains, which we will refer to hereafter as TF-ARMs.

**Figure 3.**
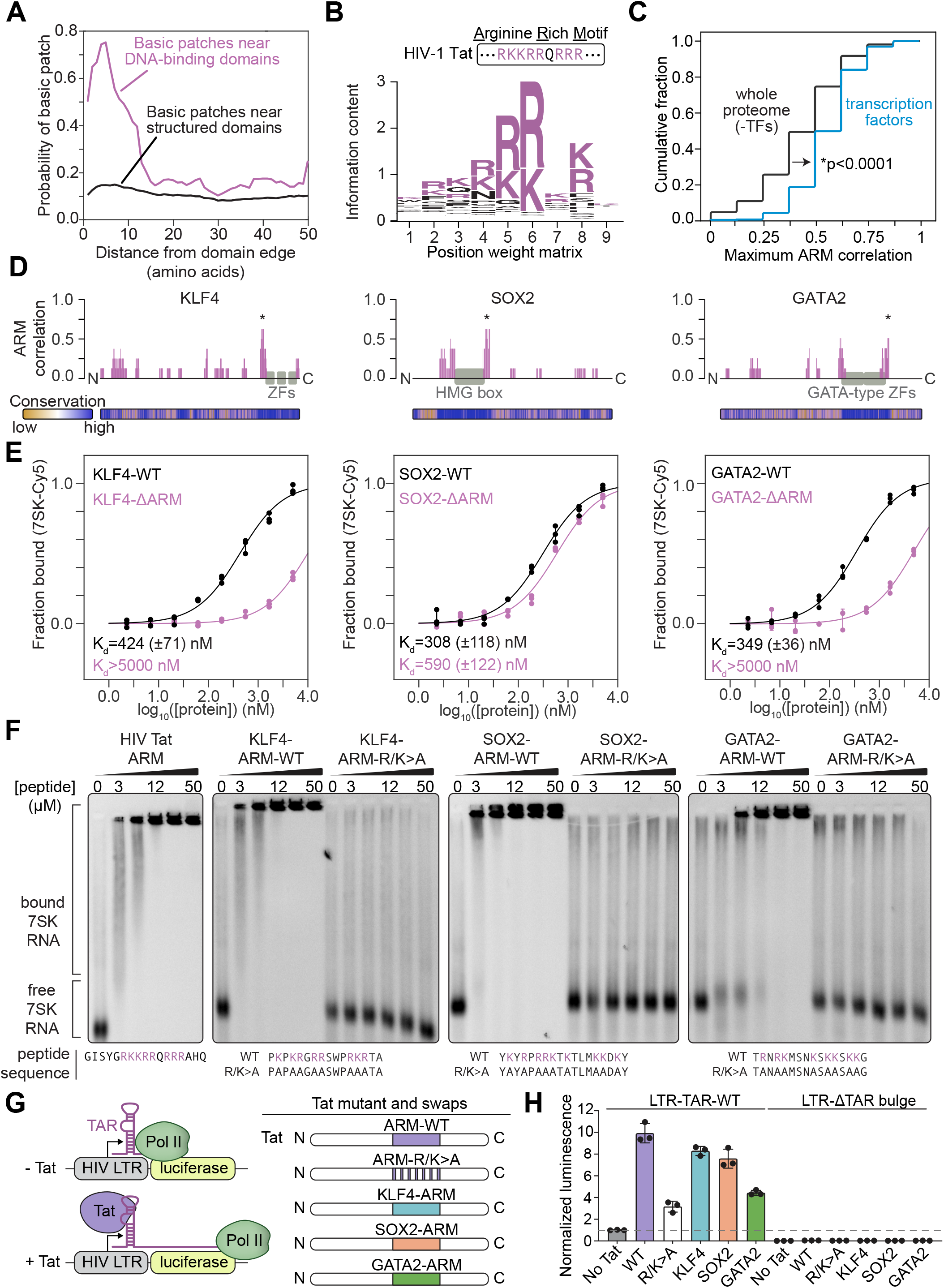
An arginine-rich domain in transcription factors. **(A)** Plot depicting the probability of a basic patch as a function of the distance from either DNA-binding domains (magenta) or all other annotated structured domains (black). **(B)** Sequence logo derived from a position-weight matrix generated from the basic patches of TFs. **(C)** Cumulative distribution plot of maximum cross-correlation scores between proteins and the Tat ARM (*p < 0.0001, Mann Whitney U test) for the whole proteome excluding TFs (black line) or TFs alone (blue line). **(D)** Diagram of select TFs and their cross-correlation to the Tat ARM across a sliding window (*maximum scoring ARM-like region). Evolutionary conservation as calculated by ConSurf (Methods) is provided as a heatmap below the protein diagram. **(E)** Fraction bound RNA with increasing protein concentration for wildtype (WT) or deletion (ΔARM) TFs (KLF4 WT vs ΔARM: p=0.017; SOX2 WT vs ΔARM: p=0.0012; GATA2 WT vs ΔARM: p=0.018). **(F)** Gel shift assay for synthesized peptides encoding wildtype or R/K>A mutations of TF-ARMs. **(G)** Experimental scheme for Tat transactivation and replacing the Tat ARM with TF-ARMs. **(H)** Bar plots depicting the normalized luminescence values for the Tat transactivation assay with or without the TAR RNA bulge. Values are normalized to the control condition (p_adj_<0.0001 for Tat RK>A compared to No Tat, WT Tat, KLF4, SOX2, and all conditions with TAR deletion; p_adj_ = 0.0086 for Tat RK>A compared to GATA2, Sidak multiple comparison test).

To investigate whether TF-ARMs are necessary for RNA binding, we purified wild-type and deletion mutant versions of KLF4, SOX2 and GATA2 and compared their RNA binding affinities. The 7SK RNA was used in this assay because it is one of a number of RNA species known to be bound by HIV Tat (Pham et al., 2018). RNA binding by the ARM-deleted proteins was substantially reduced (Figure 3E). To determine if the TF-ARMs are sufficient for RNA binding, peptides containing the HIV Tat ARM and TF-ARMs were synthesized and their ability to bind 7SK RNA was investigated using an electrophoretic mobility shift assay (EMSA). The results showed that all the TF-ARM peptides can bind 7SK RNA, as did the control HIV Tat ARM peptide (Figure 3F). This binding was dependent on arginine and lysine residues within the TF-ARMs (Figure 3F), as has been previously demonstrated for the Tat ARM (Calnan et al., 1991a; Pham et al., 2018). These results indicate that TF-ARMs are necessary and sufficient for RNA-binding.

We considered the possibility that the TF-ARM also contributes to DNA-binding. Synthesized peptides of the SOX2 and KLF4 ARMs were tested for binding to either DNA or RNA. The results show that both ARMs bind RNA with greater affinity compared to DNA (Figures S4A and S4B). Full-length wildtype and ARM-deleted SOX2 and KLF4 were also tested for binding to motif-containing DNA. The results show that deletion of the SOX2 ARM did not affect DNA-binding (Figure S4C). Deletion of the KLF4 ARM did affect DNA-binding (Figures S4D), although not to the extent that it affected RNA-binding (Figure 3E). It thus appears possible that some TF-ARMs can contribute to DNA-binding to some extent whereas others do not.

Having found that TF-ARMs bind to RNA in vitro in assays with purified components, we next asked whether TF-ARMs bind RNA in the more complex environment of the cell. To investigate this, we analyzed the RBR-ID data (Figure 1B-E), which can provide spatial information on the regions of proteins that bind RNA in cells. If TF-ARMs were binding to RNA in cells, then we would expect an enrichment of RBR-ID^+^ peptides overlapping or adjacent to the TF-ARMs. Global analysis of RBR-ID^+^ peptides in human K562 cells, as well as inspection of RBR-ID^+^ peptides for individual TFs, confirmed that this was the case (Figure S5). These results provide evidence that ARM-like regions in TFs bind to RNA in cells.

To investigate if TF-ARMs could function similarly to the Tat ARM in cells, we tested whether TF-ARMs could replace the Tat ARM in a classical Tat transactivation assay (Calnan et al., 1991a). In this assay, the HIV-1 5’ long terminal repeat (LTR) is placed upstream of a luciferase reporter gene. Transcription of the LTR generates an RNA stem loop structure called the Trans-activation Response (TAR), and HIV Tat binds to the TAR RNA to stimulate expression of the reporter gene (Jakobovits et al., 1988) (Figure 3G). We confirmed that expression of full-length Tat stimulates luciferase expression, and that mutation of the lysines and arginines in the Tat ARM reduces this activity (Figure 3H). Replacing the Tat ARM with the TF-ARMs of KLF4, SOX2, or GATA2 rescued the loss of the Tat ARM (Figure 3H). In all cases, activation was dependent on the TAR RNA bulge structure, which is required for Tat binding (Jakobovits et al., 1988) (Figure 3H). These results indicate that the TF-ARMs can perform the functions described for the Tat ARM and activate gene expression in an RNA-dependent manner.

### A role for TF RNA-binding regions in TF nuclear dynamics

TFs are thought to engage their enhancer and promoter DNA-binding sites through search processes that involve dynamic interactions with diverse components of chromatin, and we wondered if RNA-binding, enabled by TF-ARMs, contributes to TF interaction with chromatin. As an initial test of this hypothesis, we investigated whether TF-ARMs contributed to TF association with chromatin by measuring the relative levels of TFs in chromatin and nucleoplasmic fractions from ES cells containing HA-tagged TFs with wild-type and mutant ARMs. For both KLF4 and SOX2, deletion of the ARM reduced the chromatin enrichment of these TFs (Figure S6A and S6B). If RNA molecules were contributing to TF association with chromatin, then treatment of the extracts with RNase would be expected to reduce the association of TFs with chromatin; indeed, such treatment led to a loss of TFs in the chromatin fraction and an increase in the nucleoplasm (Figure S6C and S6D). These results are consistent with the idea that TF-RNA interactions can contribute to the association of TFs with chromatin.

Single molecule image analysis of TF dynamics in cells indicates that TFs conduct a highly dynamic search for their binding sites in chromatin (Chen et al., 2014; Nguyen et al., 2021). The tracking data can be fit to a three-state model, where TFs are interpreted to be immobile (potentially DNA-bound), subdiffusive (potentially interacting with chromatin components) and freely diffusing (Garcia et al., 2021a, 2021b). If TFs interact with chromatin-associated RNA through their ARMs, then we might expect that mutation of their ARMs would reduce the portion of TF molecules in the immobile and sub-diffusive states. To test this, we conducted single-molecule tracking experiments with murine embryonic stem cell (mESC) or human K562 leukemia lines that enable inducible expression of Halo-tagged wildtype or ARM-mutant TFs. For these experiments, we chose the TFs SOX2, KLF4, GATA2, and RUNX1 because of their prominent roles in mES or hematopoietic cells (Orkin and Zon, 2008; Young, 2011) and our earlier characterization of their RNA-binding regions (Figure 3). As a control, we included the deletion of an ARM-like region from CTCF that overlaps the previously described RNA-binding region (RBR) (Saldaña-Meyer et al., 2014), which was previously shown to reduce both the immobile and subdiffusive fractions of CTCF (Hansen et al., 2020). Single-molecule imaging data was fit to a three-state model (immobile, subdiffusive, and freely diffusing, Methods)(Figure 4A). Inspection of single-molecule traces for wildtype and ARM-mutant TFs (Figure 4B and Figure S7A), as well as global quantification across replicates (Figure 4C) showed that deletion of the ARM-like domains in TFs reduces the fraction of molecules in both the immobile and subdiffusive fractions, while increasing the fraction of freely diffusing molecules. The observed effects on TF mobility were consistent across different expression levels of the TFs, suggesting that the results reflect TF behavior rather than effects of over-expression (Figures S7B and S7C). These results suggest that TF-ARMs enhance the timeframe in which TFs are associated with chromatin.

**Figure 4.**
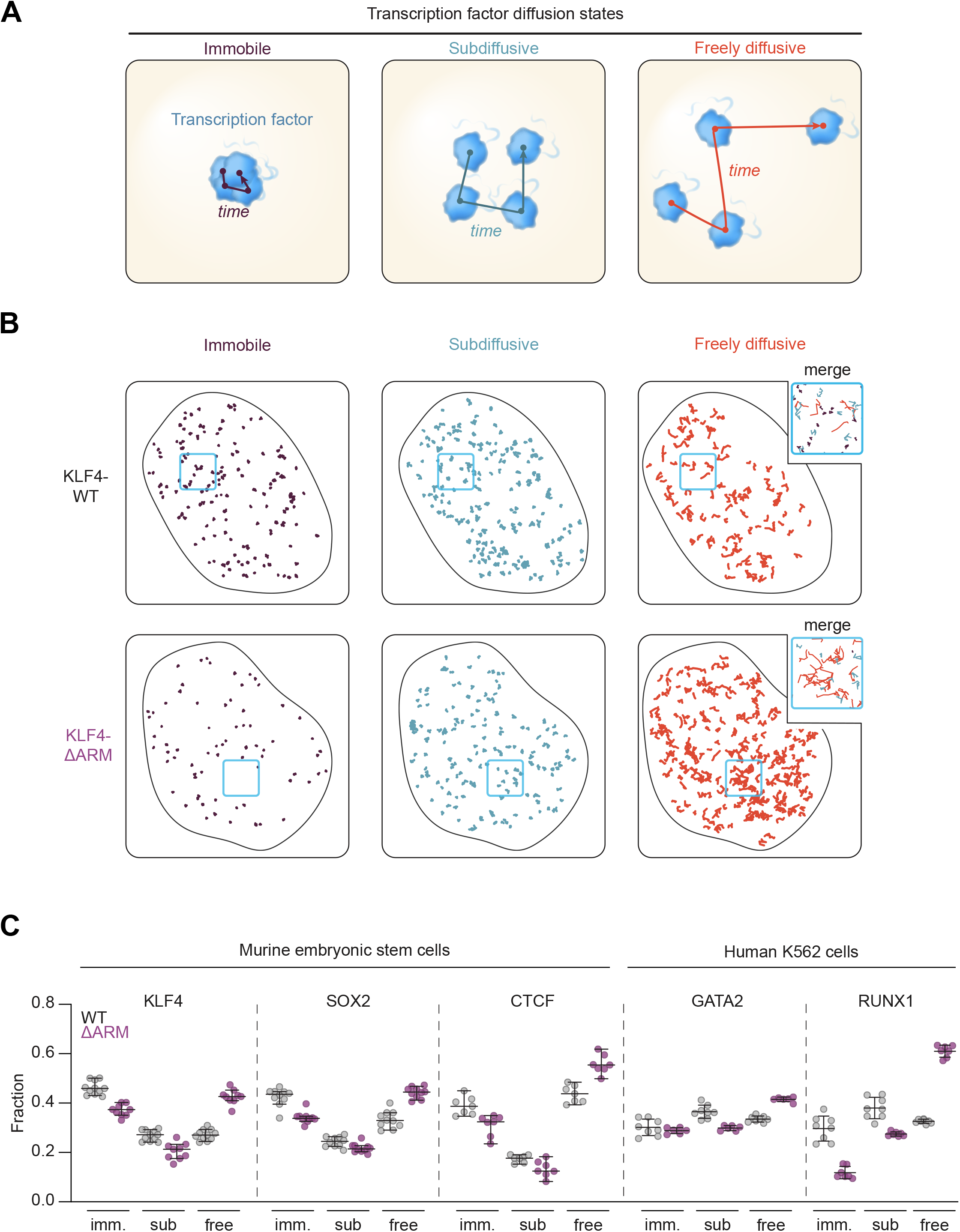
A role for TF RNA-binding regions in TF nuclear dynamics. **(A)** Cartoon depicting a 3-state model of TF diffusion. **(B)** Example of single nuclei single-molecule tracking traces for KLF4-WT and KLF4-ARM deletion. The traces are separated by their associated diffusion coefficient (D_imm_: <0.04 μm^2^s^-1^; D_sub_: 0.04-0.2 μm^2^s^-1^; D_free_: >0.2 μm^2^s^-1^). For each nucleus, 500 randomly sampled traces are shown. **(C)** Dot plot depicting the fraction of traces in the immobile, subdiffusive, or freely diffusing states. Each marker represents an independent imaging field (comparing WT and ARM-deletion, p<0.0001 for KLF4^free^, SOX2^free^, CTCF^free^, GATA2^free^, RUNX1^free^, KLF4^sub^, GATA2^sub^, RUNX1^sub^, KLF4^imm^, SOX2^imm^, RUNX1^imm^ ; p=0.0094 for SOX2^sub^; p=0.0101 for CTCF^sub^, p=0.0034 for CTCF^imm^, p=0.38 for GATA2^imm^, two-tailed Student’s t-test; error bars depict 95% C.I.).

### TF-ARMs are essential for normal development and disrupted in disease

Transcription factors are fundamental controllers of cell-type specific gene expression programs during development, so we next asked whether the TF-ARMs contribute to the factor’s role in normal development in vivo. For this purpose, we turned to the zebrafish, which has served as a valuable model system to study and perturb vertebrate development. Previous study showed that knockdown of zebrafish *sox2* by injection of antisense morpholinos at the one-cell stage led to growth defects and embryonic lethality, which could be rescued by co-injection with messenger RNA (mRNA) encoding human SOX2 (Pavlou et al., 2014). Using this system, we injected zebrafish with the *sox2* morpholino while co-injecting mRNA encoding either wildtype or ARM-mutant human SOX2 (Figure 5A), which reduced RNA but not DNA binding in vitro (Figure 3E, and Figure S4C). Embryos were scored at 48 hours post-fertilization for growth defects by the length of the anterior-posterior axis compared to embryos injected with a non-targeting control morpholino (Figure 5B). Whereas wildtype human SOX2 could partially rescue the growth defect induced by *sox2* knockdown, ARM-mutant SOX2 was unable to do so (Figure 5C). These results indicate that TF-ARMs contribute to proper development.

**Figure 5.**
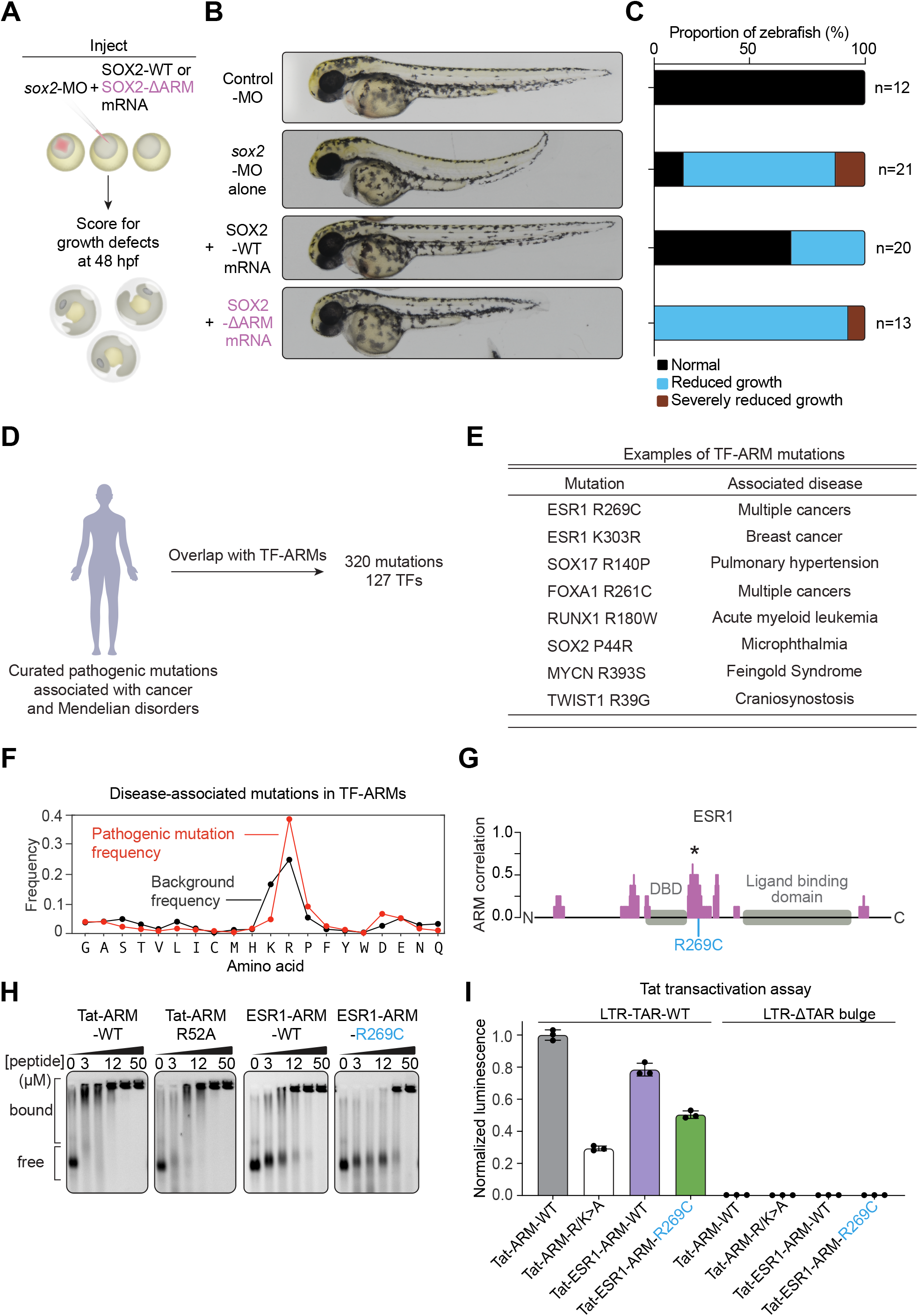
TF-ARMs are essential for normal development and disrupted in disease. **(A)** Experimental scheme for injection of zebrafish embryos with morpholinos and rescue by co-injection with mRNA (hpf = hours post-fertilization). **(B)** Representative images of injected zebrafish embryos at 48 hpf. **(C)** Scoring of zebrafish anterior-posterior axis growth. **(D)** The landscape of mutations in TF-ARMs associated with human disease **(E)** Examples of disease-associated mutations in TF-ARMs. **(F)** Line plot of the frequency of mutations (red) or background frequency (black) for amino acids in TF-ARMs (*p* = 5.2 × 10^−8^ for enrichment of mutations in arginine, one-side binomial test with Benjamini-Hochberg correction). **(G)** Representation of the ESR1 protein and its correlation to the Tat ARM (*Maximum scoring ARM-like region). The selected mutation is provided in blue. **(H)** Gel shift assay with synthesized peptides and 7SK RNA for Tat-ARM-WT, Tat-ARM-R52A, ESR1-ARM-WT, and ESR1-ARM-R269C. **(I)** Tat transactivation reporter assay with wildtype or mutant versions of Tat and ESR1 and a version of the reporter without the Tat-binding TAR bulge. Values were normalized to the Tat-WT condition.

The presence of ARMs in most TFs, and evidence that they can contribute to TF function in a developmental system, prompted us to investigate whether pathological mutations occur in these sequences in human disease. Analysis of curated datasets of pathogenic mutations revealed hundreds of disease-associated missense mutations in TF-ARMs (Figure 5D, Methods). These mutations are associated with both germline and somatic disorders, including multiple cancers and developmental syndromes, that affect a range of tissue types (Figure 5E). Variants that mutate arginine residues were the most enriched compared to the other amino acid residues in ARMs, which is consistent with their importance in RNA-binding (Figure 5F) (Calnan et al., 1991b). To confirm that such mutations could affect RNA binding, we selected for further study the estrogen receptor (ESR1) R269C mutation (Figure 5G), which is found in multiple cancers and is particularly enriched in a subset of patients with pancreatic cancer (Boldes et al., 2020). An EMSA assay showed that RNA binding was reduced with an ESR1 ARM peptide containing the R269C mutation (Figure 5H). Furthermore, when the Tat ARM was replaced with wildtype and mutant versions of the ESR1 ARM in the Tat transactivation assay, the mutation caused reduced reporter expression compared to wildtype (Figure 5I). These results support the hypothesis that disease-associated mutations in TF-ARMs can disrupt TF RNA binding.

## Discussion

The canonical view of transcription factors is that they guide the transcription apparatus to genes and control transcriptional output through the concerted function of domains that bind DNA and protein molecules (Cramer, 2019; Keegan et al., 1986; Lambert et al., 2018; Tjian and Maniatis, 1994). The evidence presented here suggests that many transcription factors also harbor RNA-binding domains that contribute to gene regulation (Figure 6). Given the large portion of TFs that showed evidence of RNA interaction in cells and the presence of an ARM-like sequence in nearly 80% of TFs, it is possible that the majority of TFs engage in RNA binding.

**Figure 6.**
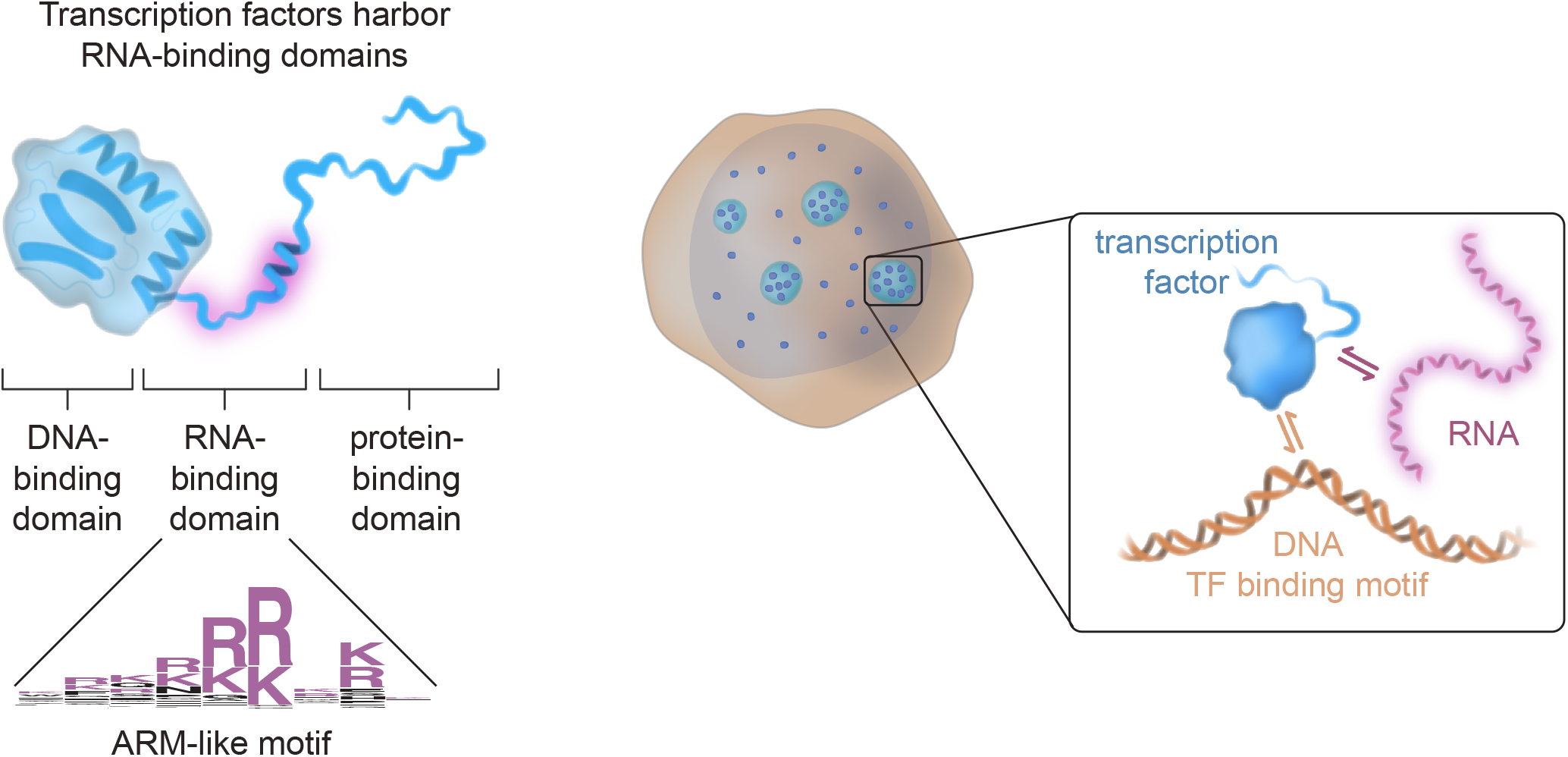
Transcription factors harbor functional RNA-binding domains. A model depiction of a previously unrecognized RNA-binding domain in a large fraction of transcription factors and its role in TF function.

RNA molecules are pervasive components of active transcriptional regulatory loci (Asimi et al., 2022; Henninger et al., 2021; Kaikkonen and Adelman, 2018; Seila et al., 2008; Sharp et al., 2022) and have been implicated in the formation and regulation of spatial compartments (Quinodoz et al., 2021). The noncoding RNAs produced from enhancers and promoters are known to affect gene expression (Kaikkonen and Adelman, 2018), and plausible mechanisms by which these RNA species could influence gene regulation have been proposed to include binding to cofactors and chromatin regulators (Bose et al., 2017; Lai et al., 2013; Long et al., 2017), and electrostatic regulation of condensate compartments (Henninger et al., 2021). The evidence that TFs bind RNA suggests additional functions for RNA molecules at enhancers and promoters. These RNA molecules serve to enhance the recruitment and dynamic interaction of TFs with active regulatory DNA loci.

The observation that many TFs can bind DNA, RNA and protein molecules offers new opportunities to further advance our understanding of gene regulation and its dysregulation in disease. Knowledge that TFs can interact with both DNA and RNA molecules may help with efforts to decipher the “code” by which multiple TFs collectively bind to specific regulatory regions of the genome and inspire novel hypotheses that may provide additional insight into gene regulatory mechanisms. It might also provide new clues to the pathogenic mechanisms that accompany GWAS variants in enhancers, where those variations occur in both DNA and RNA.

## Acknowledgements

We are grateful to Phillip Sharp, Amy Gladfelter, Seychelle Vos, and Ibrahim Cissé for discussions regarding HIV Tat, RNA-binding proteins, RNA interactions, and single-molecule imaging. We thank L.D. Lavis (HHMI, Janelia) for the gift of the Halo Tag-(PA)-JF549 dyes. This work was supported by NIH grants GM144283 (R.A.Y), CA155258 (R.A.Y.), NSF PHY2044895 (R.A.Y.), F32CA254216 (J.E.H.), GM138788 (R.B.), GM127408 (R.B.), R01 HL144780-01 (L.I.Z.), R24 OD017870-01 (L.I.Z), CA251062 (S.F.B.) and NSF Graduate Research Fellowship 1745302 (K.J.O.).

## Author contributions

Conceptualization: O.O., J.E.H., R.A.Y.

Methodology: O.O., J.E.H., R.W.T., R.B.

Software: J.E.H., K.J.O., R.W.T., M.M.Z., S.F.B., S.W.H.

Formal analysis: O.O., J.E.H.

Investigation: O.O., J.E.H., R.W.T., M.M.Z., H.E., K.J.O., S.W.H., S.F.B., R.L., A.V., N.M.H., A.L.R.

Resources: O.O., J.E.H., H.E., N.M.H.

Writing – original draft: O.O., J.E.H., R.A.Y.

Visualization: J.E.H.

Supervision: O.O., J.E.H., R.A.Y., R.B., L.I.Z., T.I.L.

Funding acquisition: R.A.Y., R.B., L.I.Z.

## Declaration of interests

R.A.Y. is a founder and shareholder of Syros Pharmaceuticals, Camp4 Therapeutics, Omega Therapeutics, and Dewpoint Therapeutics. O.O. and J.E.H. are consultants at Camp4 Therapeutics. L.I.Z. is a founder and stockholder of Fate Therapeutics, Camp4 Therapeutics, Amagma Therapeutics, Scholar Rock, and Branch Biosciences. L.I.Z. is a consultant for Celularity and Cellarity.

## Supplementary figure legends

**Figure S1.**
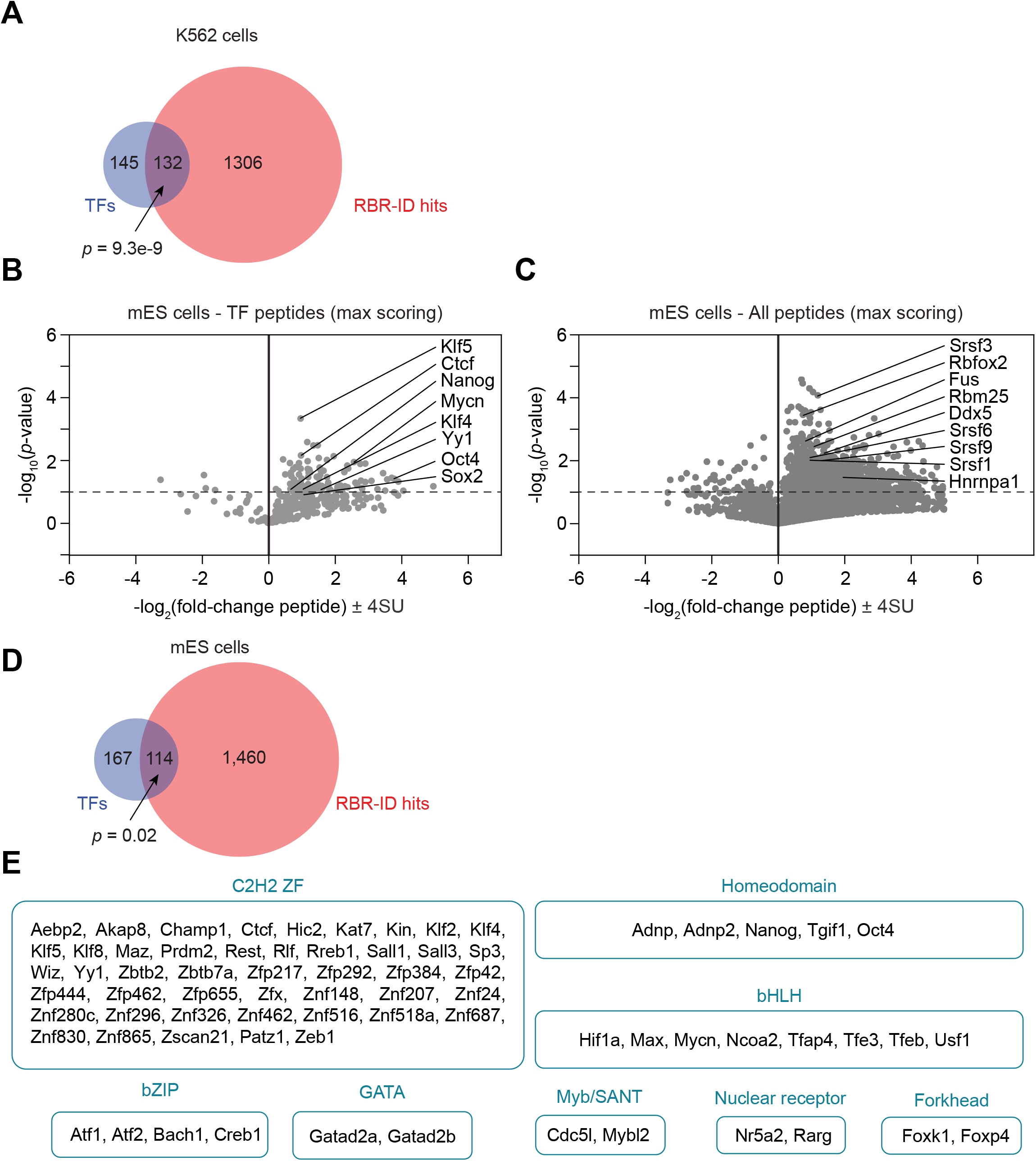
Transcription factor binding to RNA in murine embryonic stem cells (Related to Figure 1). **(A)** Venn diagram depicting overlap of RBR+ protein hits and TFs for K562 cells (p=9.3e-9, Fisher’s exact test). **(B)** Volcano plot of TF peptides in RBR-ID for murine embryonic stem cells with select highlighted TFs (dotted line at p=0.10). Each marker represents the peptide with maximum RBR-ID score for each protein. **(C)** Volcano plot of all detected peptides in RBR-ID for murine embryonic stem cells with select highlighted RBPs (dotted line at p=0.10). Each marker represents the peptide with maximum RBR-ID score for each protein. **(D)** Venn diagram depicting overlap of RBR+ protein hits and TFs for mES cells (p=0.02, Fisher’s exact test). **(E)** List of RBRID^+^ TFs (p<0.10, log_2_FC>0) categorized by DBD family.

**Figure S2.**
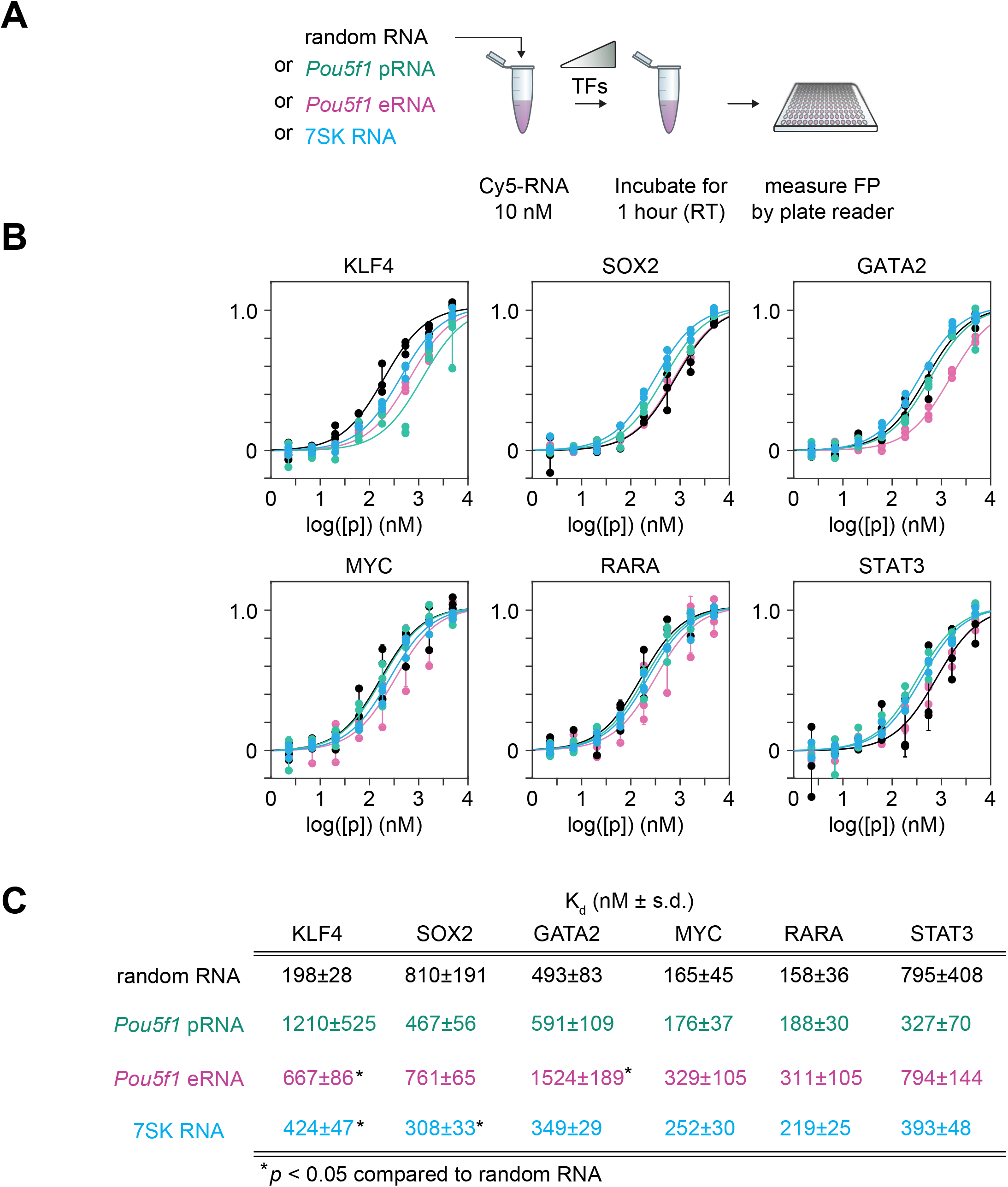
Transcription factor binding to various RNAs (Related to Figure 2). **(A)** Experimental scheme for testing TF binding affinity with multiple RNAs (eRNA = enhancer RNA, pRNA = promoter RNA). **(B)** Fraction bound RNA with increasing protein concentration for 6 TFs and 4 RNA species per TF. **(C)** Table of apparent K_d_ values for the binding assays in (B) (p-values comparing random RNA to pRNA, eRNA, and 7SK RNA respectively – KLF4: 0.06, 6.24e-6, 1.88e-4; SOX2: 0.09, 0.81, 0.013; GATA2: 0.47, 1.05e-5, 0.10; MYC: 0.84, 0.15, 0.11; RARA: 0.53, 0.17, 0.17; STAT3: 0.26, 0.99, 0.33).

**Figure S3.**
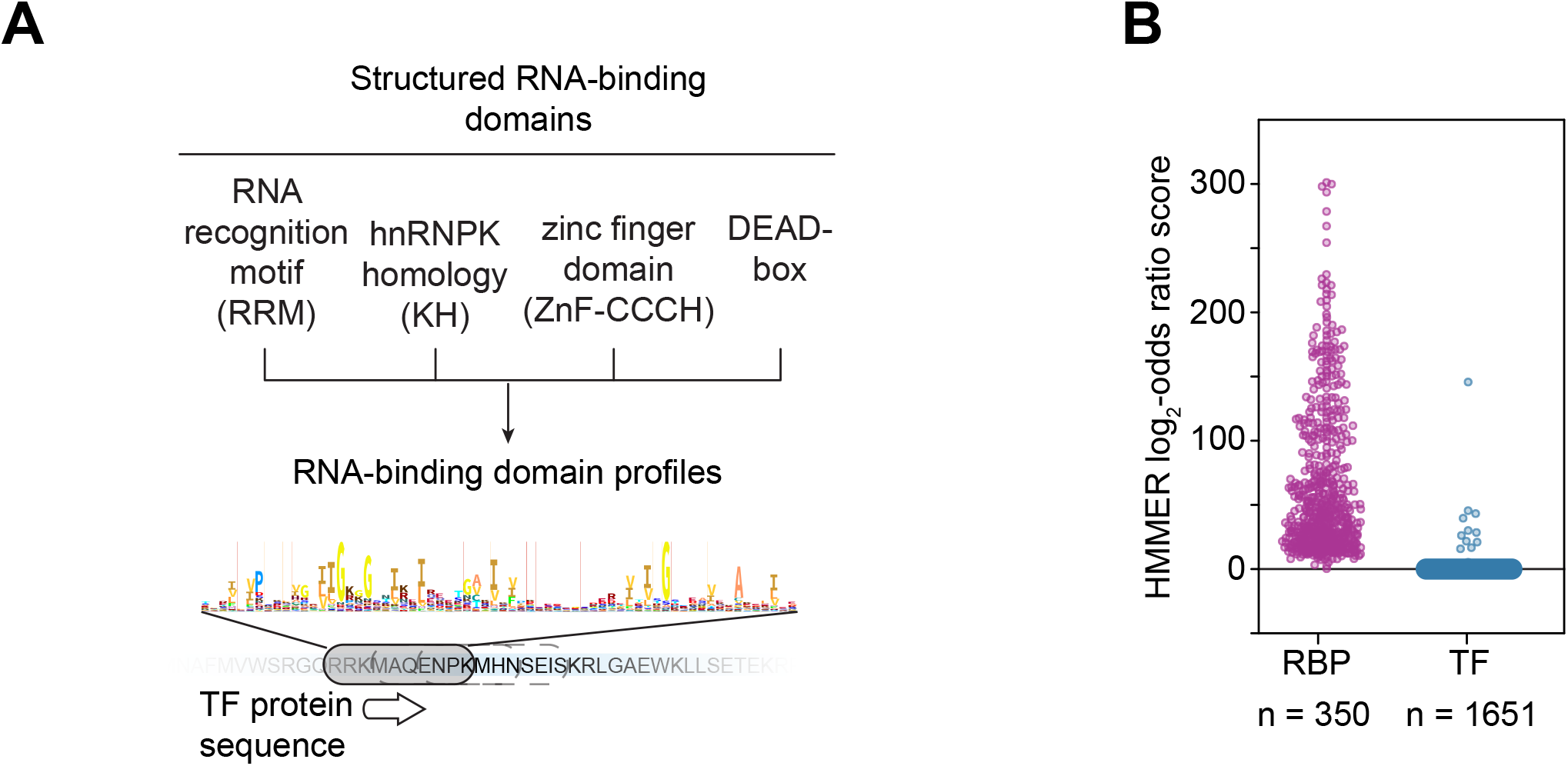
Transcription factors do not contain structured RNA-binding domains (Related to Figure 3). **(A)** Scheme to search for structured RNA-binding domain motifs in transcription factors. **(B)** Scatter plot depicting the HMMER log2-odds ratio score for the 4 most abundant RNA-binding domains (RRM, KH, ZnF-CCCH, DEAD) for select RBPs and all human TFs.

**Figure S4.**
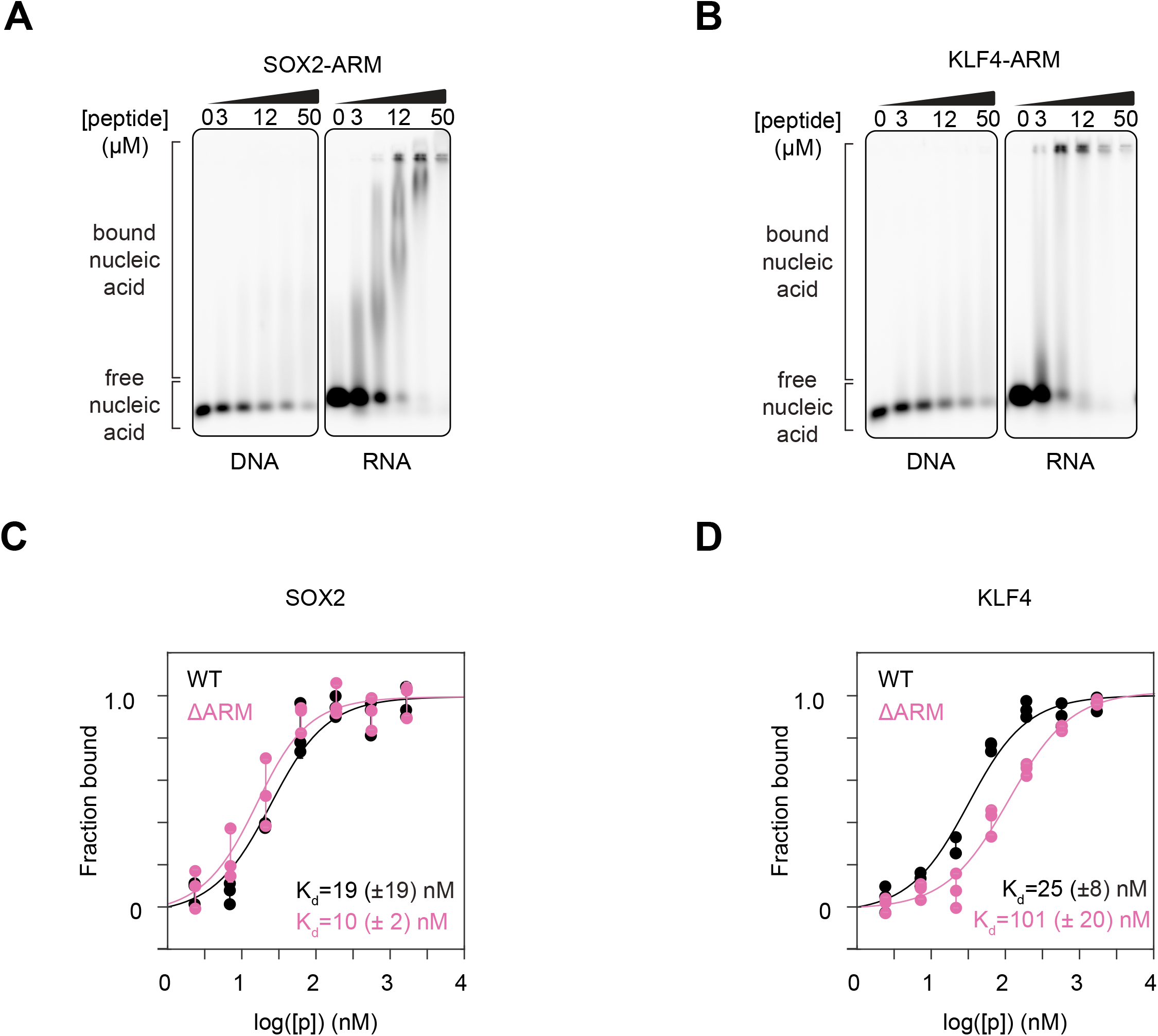
Transcription factor binding to DNA in vitro (Related to Figure 3). **(A)** Gel shift assay of the synthesized SOX2-ARM peptide with DNA or RNA **(B)** Gel shift assay of the synthesized KLF4-ARM peptide with DNA or RNA **(C)** Fraction bound motif-containing DNA with increasing protein concentration for SOX2 and **(D)** KLF4 (SOX2 WT vs ΔARM: p=0.11; KLF4 WT vs ΔARM: p=8.75e-6; error bars depict s.d.)

**Figure S5.**
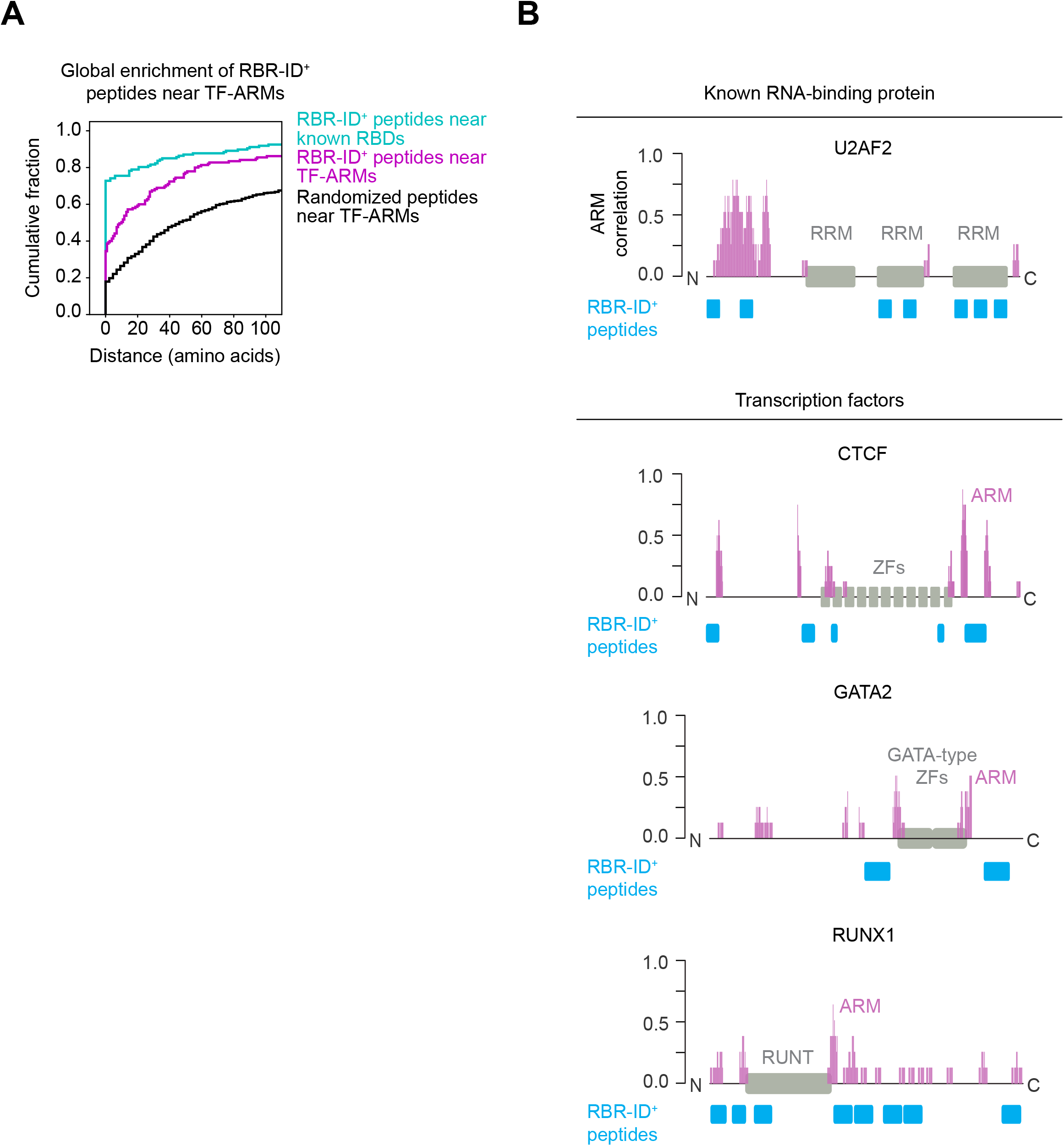
Crosslinking of TF-ARMs to RNA in cells (Related to Figure 3). **(A)** Global analysis of RBR-ID+ peptide enrichment near known RNA-binding domains (cyan), TF-ARMs (magenta), or randomized peptides near ARMs (black). **(B)** Examples of RBR-ID+ peptides for select TFs.

**Figure S6.**
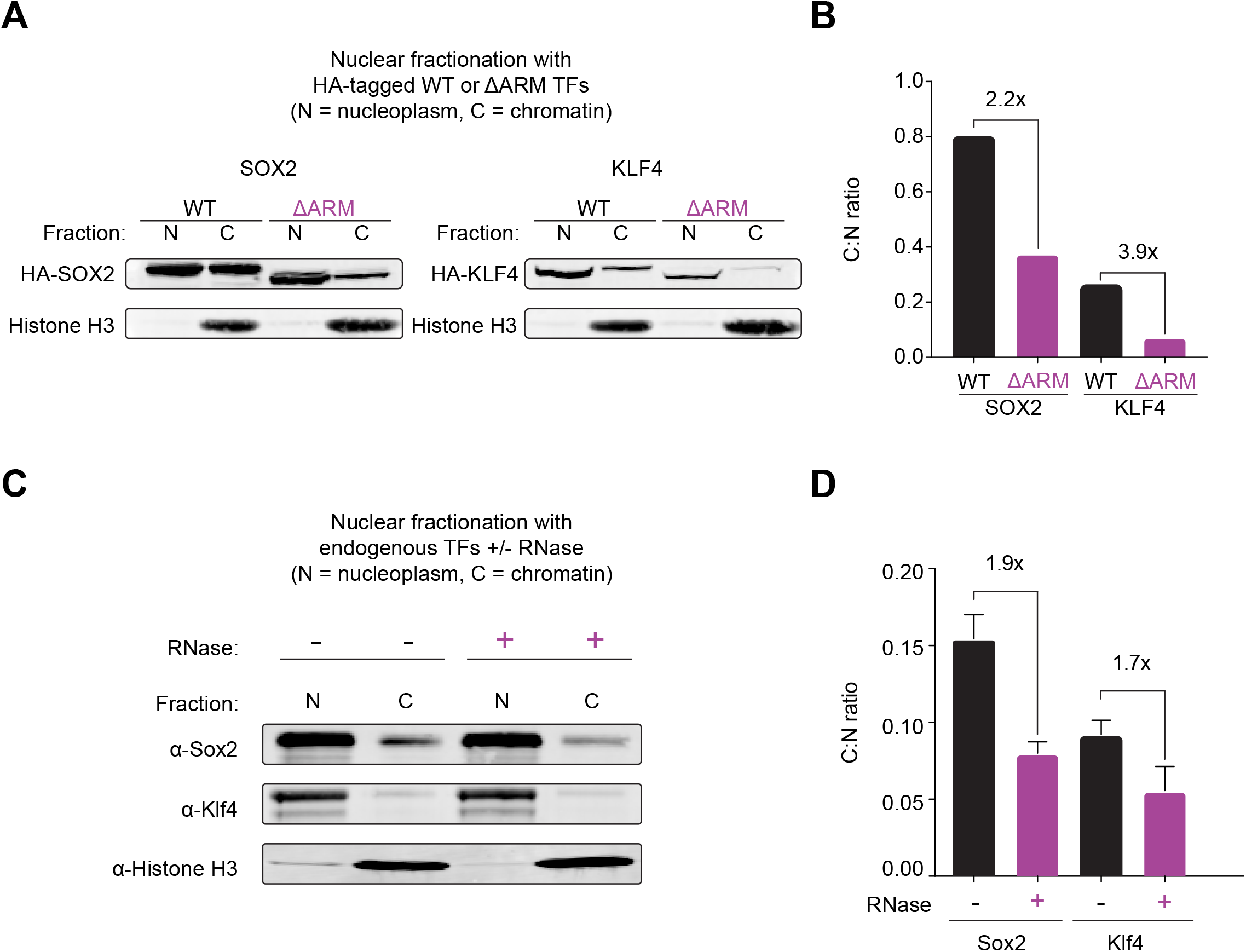
Transcription factor enrichment in sub-nuclear fractions (Related to Figure 4). **(A)** Western blot of histone H3 and HA-tagged wildtype or ARM-mutant KLF4 and SOX2 in nucleoplasmic (N) or chromatin (C) fractions. **(B)** Quantification of the relative intensity in N and C fractions of the samples in (A). **(C)** Western blot of Sox2 or Klf4 and histone H3 in nucleoplasmic (N) or chromatin (C) fractions with or without RNase treatment. **(D)** Quantification of the relative intensity in N and C fractions of the samples in **(C)**.

**Figure S7.**
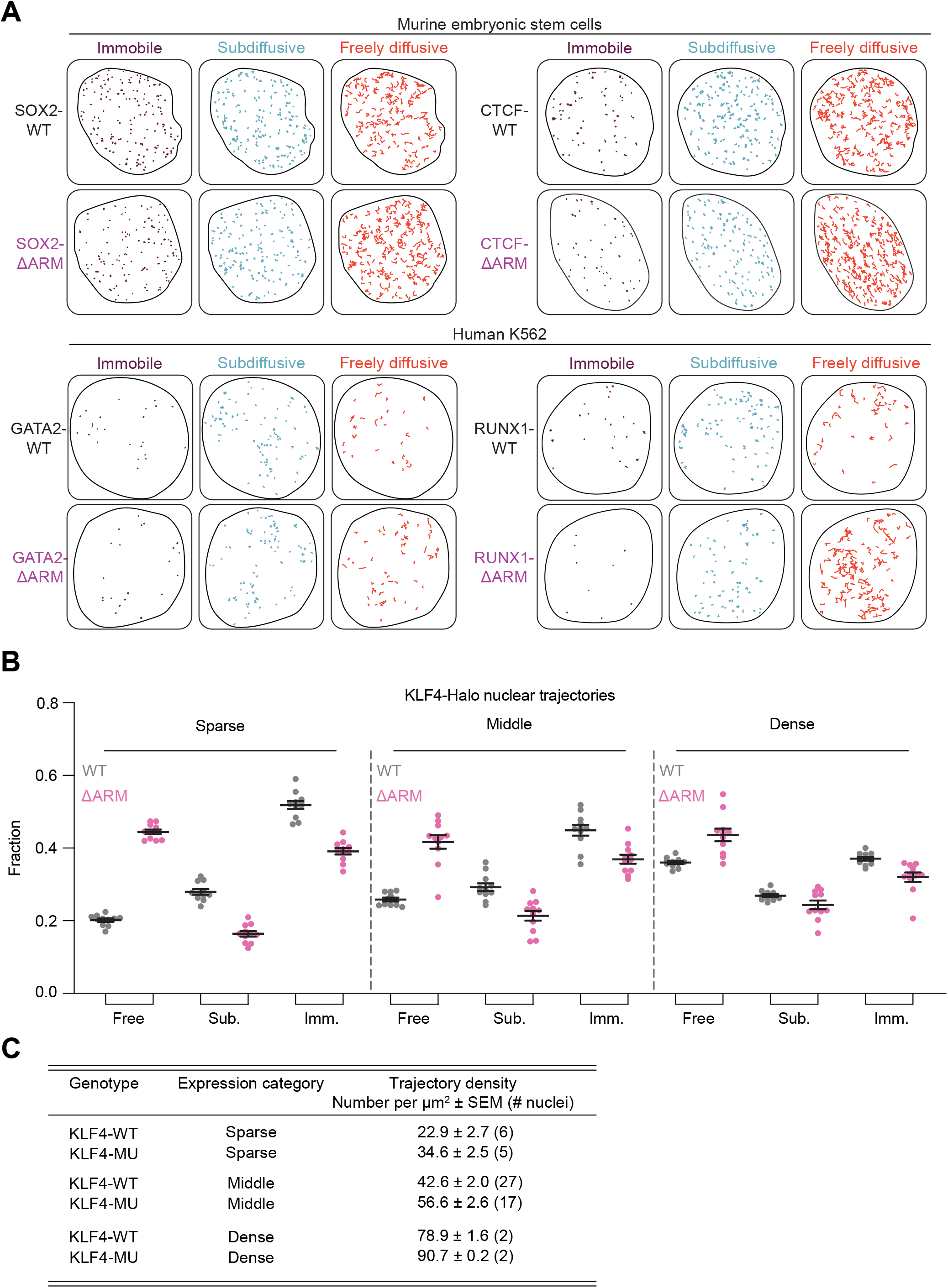
The effect of TF-ARM deletion on TF nuclear mobility (Related to Figure 4). **(A)** Example of single nuclei single-molecule tracking traces for wildtype and ARM-mutant SOX2 and CTCF in mESCs, and GATA2 and RUNX1 in K562 cells. The traces are separated by their associated diffusion coefficient (D_imm_: <0.04 μm2s-1; D_sub_: 0.04-0.2 μm2s-1; D_free_: >0.2 μm2s-1). For each nucleus, up to 500 randomly sampled traces are shown. **(B)** Fraction of traces in 3-state model across different expression levels of KLF4. **(C)** Table providing trajectory metrics across the different KLF4 expression levels.

## Supplementary information

**Table S1.** Sequences of reagents used in this study

## Methods

### Materials availability statement

All unique/stable reagents generated in this study are available from the corresponding author upon reasonable request with a completed Materials Transfer Agreement. Correspondence and material requests should be addressed to

### Data/Code Availability

The RBR-ID mass spectrometry proteomics data will be available on the ProteomeXchange Consortium via the PRIDE partner repository.

Code generated during this study will be available on Github.

### Structures of known DNA-binding domains in TFs

TF-DNA X-ray structures were obtained from the RCSB Protein Data Bank (Accession numbers: YY1 = 1UBD, MYC/MAX = 1NKP, POU2F1 = 1CQT, JUN/FOS = 1FOS). These entries were modified using ChimeraX (Goddard et al., 2018; Pettersen et al., 2021), and the effector domains, which are not included in the X-ray structures, are depicted as cartoons highlighting their dynamic and transient structure.

### RNA binding region identification (RBR-ID)

K562 cells were cultured in suspension flasks containing culture medium [RPMI-1640 medium with GlutaMAX™ (ThermoFisher Cat. 72400047) supplemented with 10% FBS (ThermoFisher Cat. 10437028), 2 mM L-glutamine (Sigma-Aldrich Cat. G7513), 50 U/mL penicillin and 50 μg/mL streptomycin]. For each biological replicate of RBR-ID, 4 million K562 cells from actively proliferating cultures were aliquoted into 2x T25 flasks. 4-thiouridine (4SU) was added to one of the two flasks for each replicate at a final concentration of 500 μM and incubated for 2 hrs at 37°C with 5% CO_2_. Cells from each flask were collected and resuspended in 600 μL 1x PBS [137 mM NaCl, 2.7 mM KCl, 10 mM Na_2_HPO_4_, 1.8 mM KH_2_PO_4_] and transferred to 6-well plates. Plates were placed on ice with their lids removed and protein–RNA complexes were crosslinked with 1 J/cm^2^ UVB (312 nm) light. Cells were lysed in Buffer A (10 mM Tris pH 7.9_4°C_, 1.5 mM MgCl_2_, 10 mM KCl, 0.5 mM DTT, 0.2 mM PMSF) with 0.2% IGEPAL CA-630 for 5 min at 4°C, then centrifuged at 2,500 g for 5 min at 4°C to pellet nuclei. Nuclei were washed 3x with 1 mL cold Buffer A (without IGEPAL) and lysed at room temperature in 100 μL denaturing lysis buffer [9 M urea, 100 mM Tris pH 8_RT_, 1x complete protease inhibitor, EDTA free (Roche Cat. 4693132001)]. Lysates were sonicated using a BioRuptor instrument (Diagenode) as follows: (energy: high, cycle: 15 sec ON, 15 sec OFF, duration: 5 min), centrifuged at 12,000 g for 10 min and supernatant was collected. Extracts were quantified using Pierce BCA assay kit (ThermoFisher Cat. 23225). 5 mM DTT was added to extracts and incubated at room temperature for one hr to reduce proteins, and then alkylated with 10 mM iodoacetamide in the dark for one hr. Samples were then diluted to 1.5 M urea with 50 mM ammonium bicarbonate and treated with 1 μL of 10,000U/μL molecular grade benzonase (Millipore Sigma Cat. E8263) and incubated at room temperature for 30 min. Sequencing grade trypsin (Promega Cat. V5117) was then added to samples at a ratio of 1:50 (trypsin:protein) by mass and incubated at room temperature for 16 hrs. The digested samples were loaded onto Hamilton C18 spin columns, washed twice with 0.1% formic acid, and eluted in 60% acetonitrile in 0.1% formic acid. Samples were dried using a speed vacuum apparatus and reconstituted in 0.1% formic acid, then measured via A_205_ quantification and diluted to 0.333 μg/μL.

For the proximity analysis in Figure S5, the nearest distance was calculated for each detected protein between RBR-ID+ peptides (p-val<0.05, log2FC<0) and either (1) TF-ARMs (cross-correlation to Tat ARM > 0.5, described below), (2) Known RNA-binding domains (RRM: IPR000504, KH: IPR004087, dsRBD: IPR014720). We required that at least 3 peptides were detected for each protein considered. As a control for the TF-ARM nearest distance analysis, the label (RBR-ID+ or RBR-ID-) of each peptide was randomly shuffled 100 times for all detected RBR-ID peptides for each protein, which provides the null distribution of the dataset.

### LC-MS/MS

Peptide samples were batch randomized and separated using a Thermo Fisher Dionex 3000 nanoLC with a binary gradient consisting of 0.1% formic acid aqueous for mobile phase A and 80% acetonitrile with 0.1% formic acid for mobile phase B. 3 μL of each sample were injected onto a Pepmax C18 trap column and washed with a 0.05% trifluoroacetic acid 2% acetonitrile loading buffer. The linear gradient was 3 minutes until switching the valve at 2% mobile phase B and increasing to 25% by 90 minutes and 45% by 120 minutes at a flow rate of 300 nL/minute. Peptides were separated on a laser-pulled 75 μm ID and 30 cm length analytical column packed with 2.4 μm C18 resin. Peptides were analyzed on a Thermo Fisher QE HF using a DIA method. The precursor scan range was a 385 to 1015 m/z window at a resolution of 60k with an automatic gain control (AGC) target of 10^6^ and a maximum inject time (MIT) of 60 ms. The subsequent product ion scans were 25 windows of 24 m/z at 30k resolution with an AGC target of 10^6^ and MIT of 60 ms and fragmentation of 27 normalized collision energy (NCE). All samples were acquired by LC-MS/MS in three technical replicates. Thermo .raw files were converted to indexed mzML format using ThermoRawFileParser utility (https://github.com/compomics/ThermoRawFileParser). To detect and quantify peptides, indexed mzML files from each set of technical replicates were searched together using Dia-NN v1.8.1 (Demichev et al., 2020) against a FASTA file of the *Homo sapiens* UniProtKB database (release 2022_02, containing Swiss-Prot + TrEMBL and alternative isoforms). Precursor and fragment m/z ranges of 300-1800 and 200-3000 were considered, respectively with peptides lengths from 6-40. Fixed and variable modifications included carbamidomethyl, N-term acetylation and methionine oxidation. A 0.01 q value cutoff was applied, and the options --peak-translation and --peak-center were enabled, while all other Dia-NN parameters were left as default.

### Bioinformatic analysis of the RBR-ID data

After removal of suspected contaminants, identified peptides were re-mapped to an updated human proteome reference (UniProtKB release 2022_02, Swiss-Prot + TrEMBL + isoforms) to reannotate matching proteins. Where multiple protein matches were identified, peptides were assigned to a single protein annotation by first defaulting to Swiss-Prot accessions, where available, then by the accession with the most matching peptides in the dataset and therefore the most likely protein group (Nesvizhskii et al., 2003). Abundances of the different charge states of the same peptide were summed, and all abundances were normalized by the median peptide intensity in each run. To assess depletion mediated by RNA crosslinking, normalized abundances for each peptide in cells treated or not with 4SU were analyzed by unpaired, two-sided Student’s *t* tests. For peptides that were missing across all 5 × 3 technical replicates in one of the treatments, Fisher’s exact tests were used comparing the frequency of peptide detection between cells treated with or without 4SU. Statistical significance was determined by adjusting *p* values from both tests using the Benjamini-Hochberg method (Hochberg and Benjamini, 1990). For mESC RBR-ID data from previous study (He et al., 2016), all peptides were re-mapped to an updated mouse reference proteome (UniProtKBrelease 2021_04) as described above while keeping original quantification and P-values. A relaxed p-value threshold (0.10) was used in the original study because it was validated to include additional RBPs (He et al., 2016). Peptides were annotated using the InterPro database (release 87, accessed 28 Feb 2022) to identify functional domains. For volcano plots, each marker represents the peptide with maximum RBR-ID score (He et al., 2016) for each protein. Transcription factors annotated in this dataset are from a previous census study (Lambert et al., 2018).

### Protein purification

To purify transcription factors, a mammalian purification system using Freestyle HEK 293F cells (gift from Sabatini lab) were used. HEK cells were grown in FreeStyle 293 Expression Medium (Gibco) on an orbital shaker. Coding sequence of desired genes were synthesized by IDT as gBlock fragments containing proper Gibson overhangs. TF-ARM deletion mutants were generated by removal of a stretch of peptide adjacent to DNA binding domains that contain ARMs. The amino acid sequences that are removed in TF-ARM mutants are shown in parentheses as follows: hsKLF4_ΔARM (aa 355-386), hsSOX2_ΔARM (aa 118-178), hsGATA2_ΔARM (aa 360-395), and hsCTCF_ΔARM (576-611). To reduce sequence complexity for gBlock synthesis, codon optimization using the IDT codon optimization tool was applied when needed. The fragments are then cloned into a mammalian expression vector containing Flag and mEGFP (N- or C-terminal) (modified from Addgene #32104) using NEBuilder HiFi DNA Assembly kit (E2611). These vectors were transiently transfected into 293F cells at a concentration of 1 million/ml with 1 μg of DNA per million cells using branched polyethylenimine (PEI) (Polysciences). 60-72 hours post-transfection, cells were resuspended in 45 ml HMSD50 buffer (20 mM HEPES pH 7.5, 5 mM MgCl2, 250 mM sucrose, 1mM DTT, 50mM NaCl, supplemented with 0.2 mM PMSF and 5 mM sodium butyrate) and incubated for 30 min at 4° C with gentle agitation. After a spin down at 3500 rpm at 4°C for 10 min, the supernatant was discarded and the pellet containing nuclei were resuspended in 35 ml of BD450 buffer (10 mM HEPES pH 7.5, 5% Glycerol, 450 mM NaCl, and protease and phosphatase inhibitors) and incubated for 30 min at 4° C with agitation. The solution was spun down at 3500 rpm at 4°C for 10 min to clear the nuclear extract. The supernatant was transferred into fresh tube and the pellet containing chromatin was passed through 18G ½ syringe 5 times. The chromatin containing lysate was spun down at 8000 rpm at 4° C for 10 min and supernatant is combined with the previously collected supernatant. Then the combined supernatants were spun down again at 8000 rpm at 4°C for 10 min to clear the lysate. 500 ul of Flag-M2 beads (Sigma) were added to the cleared lysates and incubated overnight at 4° C. The Flag-M2 beads were washed 2 times with 45 ml BD450 buffer and they were transferred into a purification column (Biorad). The beads on the column were washed 2 more times with 10 ml BD450 buffer and 5 ml Elution buffer (20 mM HEPES pH 7.5, 10% Glycerol, 300 mM NaCl). Elutions were performed by incubating the beads overnight at 4° C with 800 elution buffer and 200 ul of 5mg/ml flag peptide (Sigma). The buffer exchange (into elution buffer) and concentration of proteins were performed using spin columns (Milipore). Proteins were aliquoted and stored at -80°C.

### In vitro RNA synthesis and purification

To synthesize labeled RNA for fluorescence polarization measurements, in vitro transcription templates were generated from ssDNA oligos (for the random RNA template, Integrated DNA Technologies), gBlocks (for 7SK template, Integrated DNA Technologies), or PCR amplification of genomic DNA from V6.5 murine embryonic stem cells (for *Pou5f1* enhancer and promoter RNAs) (Henninger et al., 2021). Templates were amplified by PCR with primers containing T7 (sense) or SP6 (antisense) promoters:

T7 (added to 5’ of sense): 5’ TAATACGACTCACTATAGGG 3’

SP6 (added to 5’ of antisense): 5’ ATTTAGGTGACACTATAGAA 3’

Templates were amplified using Phusion polymerase (NEB), and the products were gel-purified using the Monarch Gel Purification Kit (NEB) following the manufacturer’s instructions and eluted in 40 μL H2O. Each template was transcribed using the MEGAscript T7 kit using 200 ng total template according to the manufacturer’s instructions. Reactions included a Cy5-labeled UTP (Enzo LifeSciences ENZ-42506) at a ratio of 1:10 labeled UTP:unlabeled UTP. The transcription reaction was incubated overnight at 37°C, and then it was incubated with 1 μL TURBO DNase (supplied in kit) for 15 minutes at 37°C. Transcribed RNA was purified by the MEGAclear Transcription Clean-Up Kit (Invitrogen) following the manufacturer’s instructions and eluting in 40 μL H2O. The RNA was diluted to 2 μM and aliquoted to limit freeze/thaw cycles. Transcribed RNA was analyzed by gel electrophoresis to verify a single band of correct size.

### Fluorescence polarization assay

To determine the binding affinity of a protein with RNA, the concentration of protein is serially diluted from 5000 nM down to 2 nM by a 3-fold dilution factor. The series of protein concentrations is then mixed with a buffer containing 10 nM cy5-labeled RNA, 10 mM Tris pH 7.5, 8% Ficoll PM70 (Sigma F2878), 0.05% NP-40 (Sigma), 150 mM NaCl, 1 mM DTT, 0.1 mg/mL non-acetylated BSA (Invitrogen AM2616), 10 μM ZnCl2. The reactions were performed in triplicates in a 20 μL reaction volume. After incubating the reactions 1 hr at room temperature, they are transferred into flat bottom black 384 well-plate (Corning 3575). Anisotropy was measured by a Tecan i-control infinite M1000 with the following parameters. Excitation Wavelength: 635 nm; Emission Wavelength: 665; Excitation/ Emission Bandwidth: 5 nm; Gain: Auto; Number of Flashes: 20; Settle Time: 200ms; G-Factor: 1. To account for instrument error, the plate was measured 3 times and the mean of the values are used in the affinity calculations. Reagents used for established RNA-binding proteins were generated previously (Guo et al., 2019) and BamHI was purchased from New England Biolabs.

To determine the binding affinity of a protein with DNA, the same buffer conditions and incubation times were used, as described above. The series of protein concentrations from 0.76-1666 nM (3-fold serial dilution) and 10 nM cy5-labeled DNA were used. The motif containing DNA sequences that have been shown to bind SOX2 (Holmes et al., 2020) and KLF4 (Sharma et al., 2021) were ordered from IDT. To prepare motif-containing DNA sequences, 50 μM of oligos with complementary sequences (one unlabeled and the other labeled with cy5) were annealed in TE+100 mM NaCl buffer by ramping down the temperature from 98°C to 4°C on a thermocycler. Then the annealed DNA fragments were diluted to appropriate concentrations with water for the assay.

Binding curves were fit to fluorescence anisotropy data via nonlinear regression with the Levenberg-Marquardt-based ‘curve_fit’ function in scipy (v. 1.7.3). Curve fitting was performed using a monovalent reversible equilibrium binding model accounting for ligand depletion, given by the equation below:

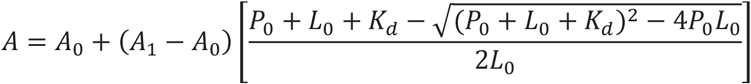

where *P*_0_ is the total protein concentration, *L*_0_ is the total ligand (RNA) concentration, and *A*_0_, *A*_1_, and *K*_*d*_ are fit parameters. The measured anisotropy value *A* for each condition was determined by first averaging raw anisotropy measurements across three subsequent reads of the same well, then averaging these values across three technical replicates from separate wells. To calculate the bound fraction of RNA, *A* values were normalized to the range between the upper and lower anisotropy asymptotes *A*_0_ and *A*_1_. Error bars were computed from the standard deviation of RNA bound fraction across three technical replicates.

### Electrophoretic mobility shift assay

To determine the binding affinity of a TF-ARM peptides (synthesized by Genscript) with 7SK RNA, the concentration of peptides was serially diluted from 50000 nM down to 3.125 nM by a 2-fold dilution factor in buffer containing 20 mM HEPES, 300 mM NaCl, and 10% Glycerol. The series of protein concentrations was then mixed 1:1 with a buffer containing an initial concentration of 20 nM Cy5-labeled RNA, 20 mM Tris pH 8.0, 5% glycerol, 0.1% NP40 (Sigma), 0.02 mM ZnCL_2_, 1 mM MgCl_2_, 2 mM DTT, and 0.2 mg/mL non-acetylated BSA (Invitrogen AM2616). For DNA-binding assays, 20 nM Cy5-labeled dsDNA or 20 nM Cy5-labeled ssRNA were used. The reactions were performed in a 20 μL reaction volume. After incubating the reactions in the dark for 1 hr at room temperature, they were loaded into a 2.5% agarose gel that is pre-run for at least 30 min at 4 °C. The samples then ran for 1.5 hr at 150V at 4 °C. The gel is imaged using Typhoon FLA95 imager with a Cy5 fluorescence module.

### Homology search for RNA-binding domains in TFs

We retrieved hidden Markov model based profiles (HMM-profiles) for RNA-binding domains corresponding to the following Pfam (Mistry et al., 2021) entries using hmmfetch from the HMMER package (hmmer.org) – RRM_1, RRM_2, RRM_3, RRM_5, RRM_7, RRM_8, RRM_9, DEAD, zf-CCCH, zf-CCCH_2, zf-CCCH_3, zf-CCCH_4, zf-CCCH_6, zf-CCCH_7, zf-CCCH_8, KH_1, KH_2, KH_4, KH_5, KH_6, KH_7, KH_8, KH_9. These domains represent the largest families of RNA-binding domains. We searched for these profiles using *hmmsearch* form the HMMER package with ‘-T 0’ as a parameter in fasta files with sequences corresponding to TFs (Lambert et al., 2018) or RNA-binding proteins (Gerstberger et al., 2014). The log2-odds ratio score from the *hmmsearch* output was plotted for RBPs with score > 0 (n=350, to provide scores that one would expect if these domains were in the protein) and for all 1651 TFs (Lambert et al., 2018). If a TF was not in the output, it was assigned a score of 0.

### Analysis of ARM-like regions in TFs

An approach based on analogous functions in localCIDER (Holehouse et al., 2017) and on a previously applied procedure (Li et al., 2020) used to map basic patches. For each TF, amino acid compositions of Lys and Arg in sliding 5-residue windows were computed. Basic patches were defined as regions of ≥ 5 consecutive residues that consisted of Lys and Arg occurring at a frequency of >0.5. This threshold was based on optimizing this approach against previously described basic patches in MECP2 (Li et al., 2020). All identified basic patches were filtered for those that occurred within predicted IDRs (*metapredict*), determined as described above. For the adjacency analysis, DNA-binding domains were defined based on domains with annotations of *DNA-binding* in Interpro (Blum et al., 2021). Probabilities of basic patch occurrence in all TFs were computed starting from the N-terminal edge of the first DNA-binding domain and moving N-terminally, or the C-terminal edge of the last DNA-binding domain and moving C-terminally. These probabilities were summed to arrive at the total probability as a function of distance from the bounds of the DNA-binding regions.

A consensus motif for bioinformatically identified basic patches (Figure 3B) was created using MEME (v. 4.11.4) (Bailey et al., 2009). Briefly, 963 basic patches found in TFs were padded by appending the 10 amino acid residues upstream and downstream of each the region. Next, a zero-order Markov model was created from 1,290 full sequences of annotated TFs using the ‘fasta_get_markov’ function to generate a background for the motif search. The TF basic patch sequences were input to the ‘MEME’ function using the TF background model, specifying a constraint to identify exactly one site per sequence, a minimum motif width of 5, a maximum motif width of 13, and defaults for the unspecified parameters.

A charge-based cross-correlation method was employed to identify ARMs in TF disordered regions similar to the HIV Tat ARM. Extensive in vitro and cellular analyses of the Tat ARM have mapped the critical residues responsible for Tat RNA-binding and HIV transactivation (Calnan et al., 1991a, 1991b). To properly function, the Tat ARM requires an arginine positioned near the motif center flanked by an enrichment of basic residues (R/K). The Tat ARM sequence “RKKRRQRRR” was digitized to the amino acid charge pattern “111110111” to create a 9-mer search kernel. A protein target sequence was created by first digitizing the sequence of the protein of interest to “1” for R/K amino acid residues and “0” otherwise, then refining the sequence by setting residues to “0” if they fell outside of disordered regions assessed through the *metapredict* package (Emenecker et al., 2021) (v. 2.2) with a disorder threshold of 0.2. The target sequence was further refined by setting all entries to “0” in 9-mer windows where no R’s were originally present. The cross-correlation between the search kernel and the target sequence was then computed using the ‘correlate’ function in scipy using the “direct” method. Maximum cross-correlations were computed as the maximum of the returned array for each protein tested. This method was applied iteratively to all sequences from the UniProt database to generate distributions for TFs and the proteome.

### Evolutionary conservation of TF-ARMs

Evolutionary conservation of human TFs was assessed using the ConSurf online server (Ashkenazy et al., 2016). TF sequences were downloaded from UniProt and run without specifying a 3D structure or MSA, with automatic detection of homologs from the “NR_PROT_DB” database. Defaults were used for all other running parameters. Amino acid conservation scores from the ConSurf GRADES output were re-normalized between 0 and 1 for each protein, such that a score of 1 corresponded to the of the most conserved amino acid in a given protein.

### HIV Tat transactivation assay

To generate the HIV LTR luciferase reporter, the HIV 5’ LTR from the pNL4-3 isolate (Genbank AF324493) was cloned into pGL3-Basic (Promega) via Gibson assembly (NEB 2X HiFi) with a HindIII-digested pGL3-Basic and a gBlock (Integrated DNA Technologies) containing the HIV 5’ LTR with compatible overhangs. A mutant version of this reporter lacking the Tat activation site (TAR RNA bulge structure) (Jakobovits et al., 1988) was also generated in a similar fashion. Mammalian expression vectors encoding Tat, an R/K>A mutant of Tat, and replacements of the Tat ARM with TF-ARMs from KLF4, SOX2, GATA2, and ESR1 were generated by Gibson assembly with a NotI-XhoI-digested pcDNA3 (Invitrogen) and gBlocks encoding these variants with compatible overhangs.

For transfections, HEK293T cells were cultured in DMEM (Gibco) supplemented with 10% fetal bovine serum (Sigma F4135), 50 U/mL penicillin and 50 μg/mL streptomycin (Life Technologies 15140163). Transfections were conducted in triplicate. 24-well plastic plates were first coated with poly-L-lysine (Sigma) for 30 minutes at 37°C, washed once with 1X PBS, and then allowed to air dry. Cells were seeded in 500 μL of media in coated wells at a density of 2×10^5^ cells per well. The next day, each well was transfected using Lipofectamine 3000 (Life Technologies) (total reaction 50 μL Optimem, 1.5 μL Lipo-3000, 0.6 μL P3000, and the appropriate volume of DNA) with 100 ng of the HIV 5’ LTR reporter vector, 150 ng of the pcDNA3 expression vector (encoding Tat or the variants), and 50 ng of a renilla luciferase plasmid (pRL-SV40, Promega) to normalize transfection efficiency. As a control, we included a pcDNA3 vector expressing LacI-mCherry (labeled as “No Tat” in Figure 3). After 6 hours of incubation, luciferase activity was quantified by the Dual Luciferase Assay kit (Promega) following the manufacturer’s instructions and a Safire II plate reader. The luminescence values were first normalized to the renilla luciferase luminescence for each well, and then all conditions were normalized to the average value of the “No Tat” control condition.

### Single-molecule tracking

#### Cell line generation

Murine embryonic stem cells were cultured in 2i/LIF media on tissue culture plates coated with 0.2% gelatin (Sigma, G1890). The 2i/LIF media contained: 960 mL DMEM/F12 (Life Technologies, 11320082), 5 mL N2 supplement (Life Technologies, 17502048; stock 100X), 10 mL B27 supplement (Life Technologies, 17504044; stock 50X), 5 mL additional L-glutamine (GIBCO 25030-081; stock 200 mM), 10 mL MEM nonessential amino acids (GIBCO 11140076; stock 100X), 10 mL penicillin-streptomycin (Life Technologies, 15140163; stock 10^4 U/mL), 333 mL BSA fraction V (GIBCO 15260037; stock 7.50%), 7 mL b-mercaptoethanol (Sigma M6250; stock 14.3 M), 100 mL LIF (Chemico, ESG1107; stock 10^7 U/mL), 100 mL PD0325901 (Stemgent, 04-0006-10; stock 10 mM), and 300 mL CHIR99021 (Stemgent, 04-0004-10; stock 10 mM). Cells were passaged by washing once with 1X PBS (Life Technologies, AM9625) and incubating with TrypLE (Life Technologies, 12604021) for 3-5 minutes, then quenched with serum-containing media made by the following recipe: 500 mL DMEM KO (GIBCO 10829-018), MEM nonessential amino acids (GIBCO 11140076; stock 100X), penicillin-streptomycin (Life Technologies, 15140163; stock 10^4 U/mL), 5 mL L-glutamine (GIBCO 25030-081; stock 100X), 4 mL b-mercaptoethanol (Sigma M6250; stock 14.3 M), 50 mL LIF (Chemico, ESG1107; stock 10^7 U/mL), and 75 mL of fetal bovine serum (Sigma, F4135). Cells were passaged every 2 days.

A piggyBac compatible base vector was assembled containing two tandem gene cassettes: (1) an insertion site downstream of a doxycycline-inducible promoter allowing for the expression of a Flag-HA-Halo-tagged ORF with SV40 NLS and bGH polyA termination sequence, and (2) the Tet-On 3G rtta element driven by the EF1a promoter that also produces hygromycin resistance via a 2A self-cleaving peptide. This base vector was generated by Gibson assembly. Plasmids encoding Halo-tagged versions of TFs (WT and ARM-deletion) were generated by Gibson assembly with BamHI-digested base vector and gBlocks (Integrated DNA Technologies) encoding the WT and ARM-deletion TFs.

To generate cell lines, 5×10^6^ mESCs per well were transfected in 6-well plates with 1 μg of the Halo-TF vector and 1 μg of the piggyBac transposase (Systems Biosciences) in serum-containing media (described above) using Lipofectamine-3000 for at least 4 hours. After transfection, the cells were passaged into 10 cm plates in 2i media containing 500 ng/mL Hygromycin-B (Gibco 10687010). After 2-4 days of selection, cells were maintained as described above.

#### Sample preparation

Cells were plated on glass bottom dishes (Cellvis D35-20-1.5-N) coated with 5 μg/ml of poly-L-ornithine (Sigma-Aldrich P4957) for 2hrs min at 37°C and with 5μg/ml of Laminin (Corning® 354232) for 2hrs-24hrs at 37°C, growing from 20% confluency in 2i for one day. Doxycycline=10ng/mL was added to dishes for 1hr, followed by adding 5nM of HaloTag-(PA) JF549 for another 3hrs. Cells were then rinsed once with PBS and washed in fresh 2i for 1hr. Dishes were refilled with 2mL prewarmed Leibovitz’s L-15 Medium, no phenol red (ThermoFisher 21083027) and brought for imaging.

#### Imaging

Cells were imaged on an inverted, widefield setup with a Nikon Eclipse Ti microscope and a 100x oil immersion objective as previously described (Henninger et al., 2021). Images were acquired with an EMCCD camera (EM gain 1000, exposure time 10ms, conjugated pixel-size on sample 160nm). A 561nm laser beam of 150mW (attenuated with 50% AOTF) was 2x expanded for a uniform illumination across around 200×200 pixel region. 10,000 frames were recorded for each ROI (including 2-4 cells), and the 405nm activation was kept very low to guarantee the molecule sparsity needed for robust reconnection.

#### Analyses

Particle trajectories were detected and reconnected with customized MATLAB code from MTT (Sergé et al., 2008). Detection settings: false-positive threshold=24, window-size 7×7pixel, and Gaussian width fitting allowed. Reconnection settings: T_off_=10ms, T_cut_=20ms, and r_max_=270nm. A collection of trajectories from each ROI were fitted to a 3-state model in Spot-on (Hansen et al., 2018). Spot-on settings: detection slice dZ=950nm, 8 delays to consider, and only first 10 jumps to consider for each trajectory. The final outputs include fractions and apparent diffusion coefficients of each state (immobile, sub-diffusive, and free, respectively). For expression dependence testing in Figure S5B, trajectories of the same genotype from different nuclei with similar trajectory density were gathered together first and resampled ten times (2,000 trajectories for each resampling) for ten independent Spot-on fittings, respectively. In this way, the accuracy of each fitting and the distributions across different conditions are comparable.

### Sub-nuclear fractionation

mESCs with exogenous expression for SOX2 and KLF4 wild type and ARM deletion mutations expressing HA tag were used for nuclei sub fractionation. To extract nuclei, cells were resuspended in 10 ml HMSD50 buffer (20 mM HEPES pH 7.5, 5 mM MgCl2, 250 mM sucrose, 1mM DTT, 50mM NaCl, supplemented with 0.2 mM PMSF and 5 mM sodium butyrate) and incubated for 30 min at 4°C with gentle agitation. After a spin down at 3500 rpm at 4°C for 10 min, the supernatant was discarded and the pellet containing nuclei were subjected to subcellular protein fractionation for nucleoplasm and chromatin fractions using the Subcellular Protein Fractionation Kit for Cultured Cells (ThermoScientific, Ref 78840) according to manufacturer’s instructions. For RNase treatment in wild type mESCs, nuclei were treated with RNase A (1:100, Thermo Fisher EN0531) and the initial 30-minute incubation at 4°C was adjusted to 20 minutes at 4°C and 10 minutes at 37°C. SDS Page was run on 12% Bis-Tris gel (Criterion XT, BioRad) and western blotting was performed on the subfractions using anti Histone H3 antibody from Abcam (ab1791) and anti HA antibody from Abcam (ab9110) with secondary antibody against Rabbit (IRDye 800CW Goat anti-rabbit LI-COR 926-32211). For wild type transcription factor detection, antibody for Sox2 (R&D Systems, MAB2018) and Klf4 (R&D Systems, AF3158) with secondary antibody anti-mouse for Sox2 (IRDye 680CW goat anti-mouse LI-COR 926-32211) and anti-goat for Klf4 (IRDye 800CW donkey anti-goat LI-COR 926-32214), were used. Fluorescence was assessed using Odyssey CLX LiCOR and quantified using ImageJ.

### Zebrafish knockdown and rescue of *sox2*

Morpholinos (MO, GeneTools) were resuspended in nuclease free water, heated to 65°C for 5 minutes, and stored at room temperature. Wildtype AB zebrafish embryos were injected into the yolk at the 1-cell stage with 7ng of *sox2*-MO (TCTTGAAAGTCTACCCCACCAGCCG) (Pavlou et al., 2014), either alone or in combination with 25 pg of human wildtype or ARM-deletion SOX2 mRNA. Messenger RNA was synthesized using the T7 mMessage mMachine (Invitrogen) kit with templates generated from gBlocks (IDT). The mRNA was purified with the MEGAclear Clean-Up Kit (Invitrogen), run on a TBE agarose gel to confirm purity and size, aliquoted, and stored at -80°C. Embryos injected with 7ng of Standard Control MO (CCTCTTACCTCAGTTACAATTTATA) were used as controls. At 48 hours post fertilization (hpf), MO injected embryos were dechorionated using forceps, anaesthetized using 0.16 mg/ml Tricaine, then visually assessed for growth impairment using a Nikon SMZ18 stereoscope with DS-Ri2 camera and NIS-Elements software. Embryos were scored based on rescue of growth impairment in the presence of wildtype or mutant sox2 mRNA.

### Overlap of pathogenic mutations in TF-ARMs

Pathogenic nonsynonmous substitution mutations were obtained from a prior dataset of pathogenic mutations that integrated multiple databases of somatic and germline variation associated with cancer and Mendelian disorders, including ClinVar (accessed January 29, 2021) and HGMD v2020.4 in hg38. Cancer variants were obtained from AACR Project GENIE v8.1 (AACR Project GENIE Consortium, 2017) and various TCGA and TARGET studies via cBioPortal (accessed January, 2021)(Banani et al., 2022). Mutations were subsetted for those affecting TF-ARMs. The frequency of the mutated wild-type amino acid across these TF-ARM mutations was compared to the frequency of the same amino acid across all TF-ARMs, and enrichment was defined as a higher pathogenic mutation frequency compared to the background amino acid frequency. Statistical significance of the enrichment was determined using a one-sided binomial test, and p-values were corrected for the multiple tests across the twenty amino acids using the Benjamini-Hochberg method.

### Statistical information

Confidence intervals for K_d_ estimates from fluorescence polarization data were computed by multiplying the standard deviation of the K_d_ curve fit parameter with the Student’s t-value corresponding to the 95% confidence interval with degrees of freedom equal to the number of data points in the concentration curve minus the number of fit parameters. Statistical comparisons between the K_d_’s of two fluorescence polarization curves (for Figure 3E, Figure S2C, and Figure S4) were assessed using a two-tailed Student’s t-test based on the standard errors of the K_d_ parameters calculated from the diagonals of the covariance matrix returned by ‘curve_fit’ in scipy, with the degrees of freedom as specified above.

The distributions of ARM correlation scores (Figure 3C) for whole proteome (-TFs) vs TFs were compared using a two-tailed Mann Whitney U test, n1=1287, n2=20238.

The Tat reporter assays were conducted on 3 biological replicates per genotype, and luminescence readings were measured in technical duplicates. Each condition was compared to the Tat R/K>A condition using a Sidak multiple comparisons test (DF = 24, t statistics were as follow: TAR-WT - WT=20.15, KLF4=15.3, SOX2=13.17, GATA2=3.805, NoTat=6.419; ΔTAR-bulge – WT=9.263, KLF4=9.319, SOX2=9.329, GATA2=9.315, Tat R/K>A=9.302, No-Tat=9.364).

For comparison of the diffusive fractions reported in Figure 4C, multiple fields of cells were imaged per genotype (KLF4-WT n=11, KLF4-ΔARM n=9, SOX2-WT n=10, SOX2-ΔARM n=9, CTCF-WT n=7, CTCF-ΔARM n=7). The diffusive fractions were compared by 2-tailed Student t-test. The data was confirmed to have equal variance via F test, and the degrees of freedom and t statistics were as follows: KLF4-free (t=13.47, df=18), SOX2-free (t=8.297, df=18), CTCF-free (t=6.044, df=12), KLF4-sub (t=5.152, df=18), SOX2-sub (2.908, df=18), CTCF-sub (t=3.051, df=12), KLF4-imm (t=7.824, df=18), SOX2-imm (t=6.203, df=18), CTCF-imm (t=3.639, df=12).

